## Supplemental figures and tables for "Nanoscale analysis of human G1 and metaphase chromatin *in situ*"

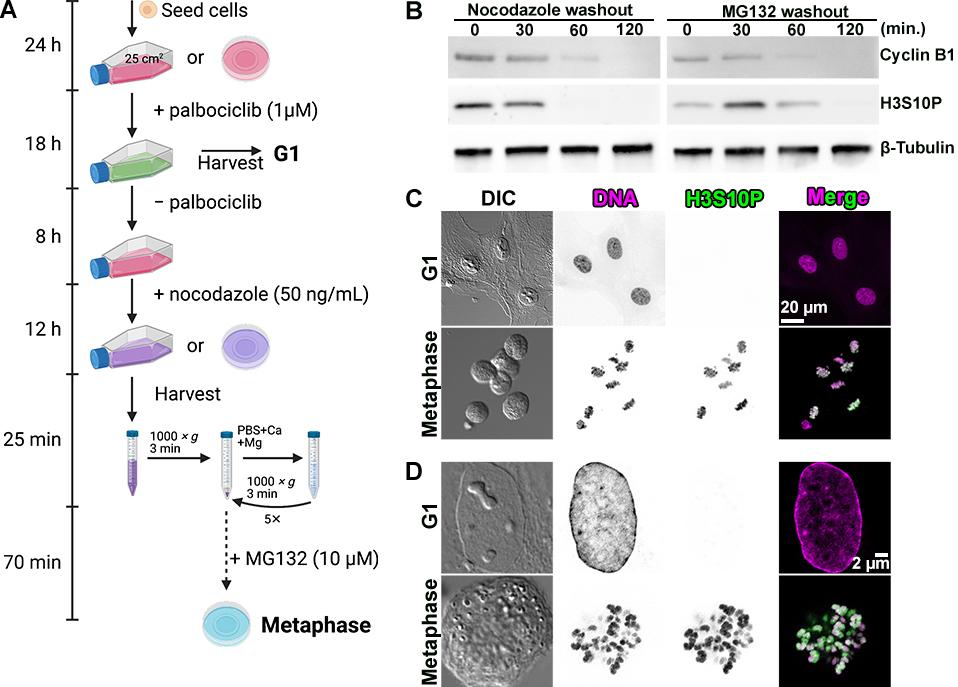

**Figure S1. Synchronization scheme.**

(A) Schematic of RPE-1 G1 and metaphase cell-cycle synchronization. Created with BioRender.com. (B) Immunoblot analysis showing decrease of mitotic markers cyclin B1 and histone H3 phosphorylated at serine 10 (H3S10P) 0 – 120 minutes following washout of nocodazole and MG132. β-Tubulin is the loading control. The uncropped blots are shown in Figure S36. (C) Differential interference contrast (DIC) and immunofluorescence images of G1 and metaphase cells. The chromatin is stained with DAPI (DNA). Immunostaining for histone H3 phosphorylated at serine 10 (H3S10P). The fluorescence contrast is inverted for better visibility. (D) Airyscan sections of a G1 and a metaphase cell, showing the localization of H3S10P in the metaphase chromosomes.

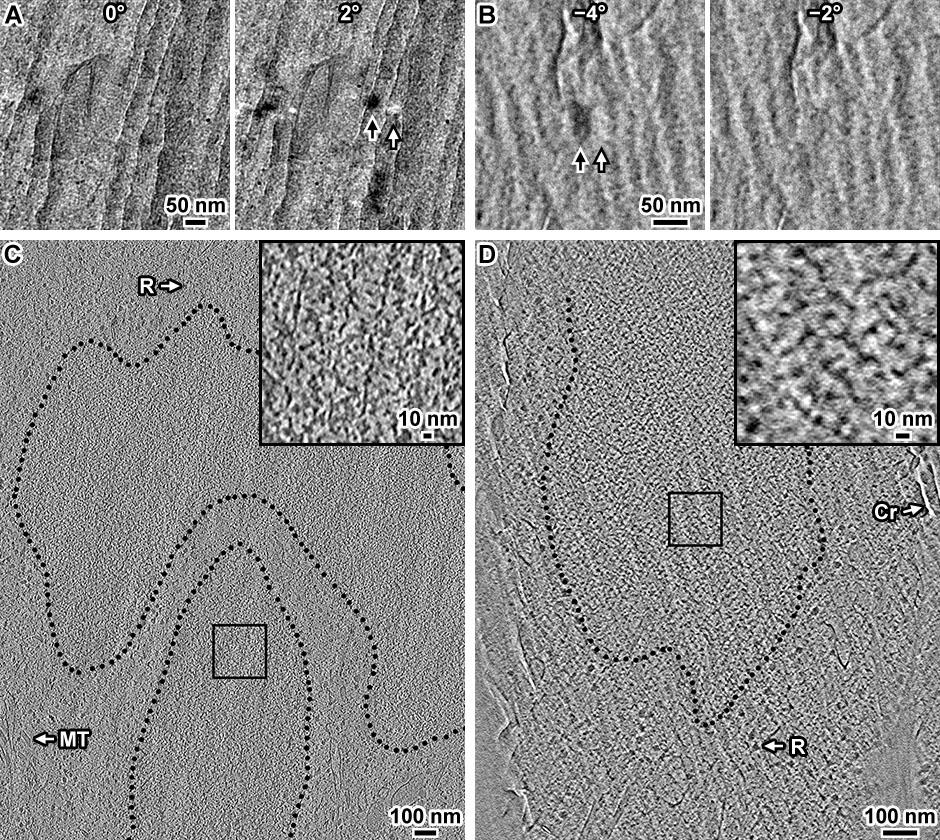

**Figure S2. Cryosections of RPE-1 cells in metaphase.**

(A and B) Pairs of tilt series images of two RPE-1 cells. In each panel, the two images correspond to a 2° difference in tilt angle. The black and white arrows indicated diffraction contrast features that arise from crystalline ice within the cellular cryosection. The linear vertical features are crevasses. (C and D) Cryotomographic slices (12 nm) of the cells in panels A and B, respective. Cytological features such as ribosomes (R) and a microtubule (MT) are indicated. (Cr) indicates a crevasse feature. Black dotted lines indicate the approximate boundary that encloses the compacted chromosomes. The insets show 4-fold enlargements of the boxed areas in the chromosomes.

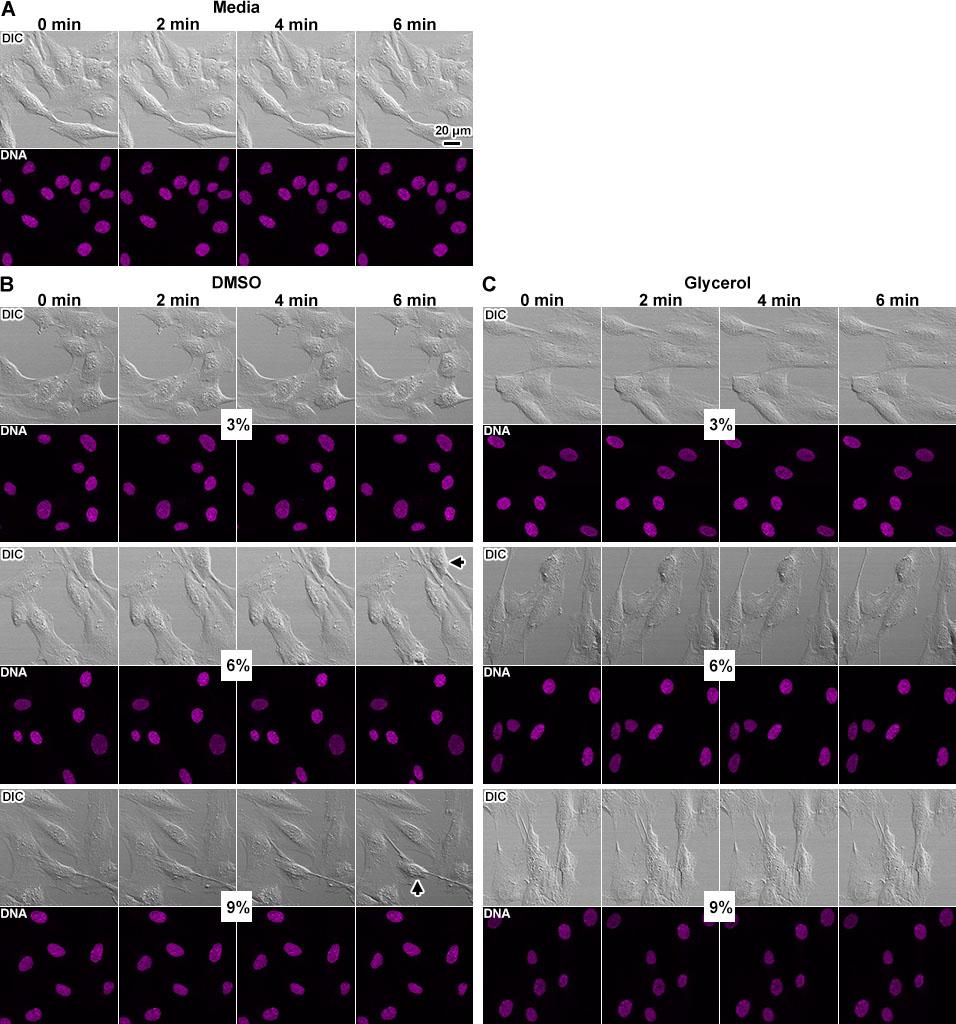

**Figure S3. Screen of cryoprotectants in live cells.**

(A) Control timelapse images of unsynchronized RPE-1 cells in complete media and stained with Hoechst 33342, imaged every 2 minutes for 6 minutes. Images were acquired in the DIC (upper row) and Hoechst 33342 (lower row) channels. The experimental delay between the addition of cryoprotectant and the acquisition of the first image is estimated to be 1 minute. Timelapse imaging was done for cells in either (B) DMSO or (C) glycerol, at three different concentrations (3%, 6%, and 9%). The short arrows in panel B indicate cells that have started to detach.

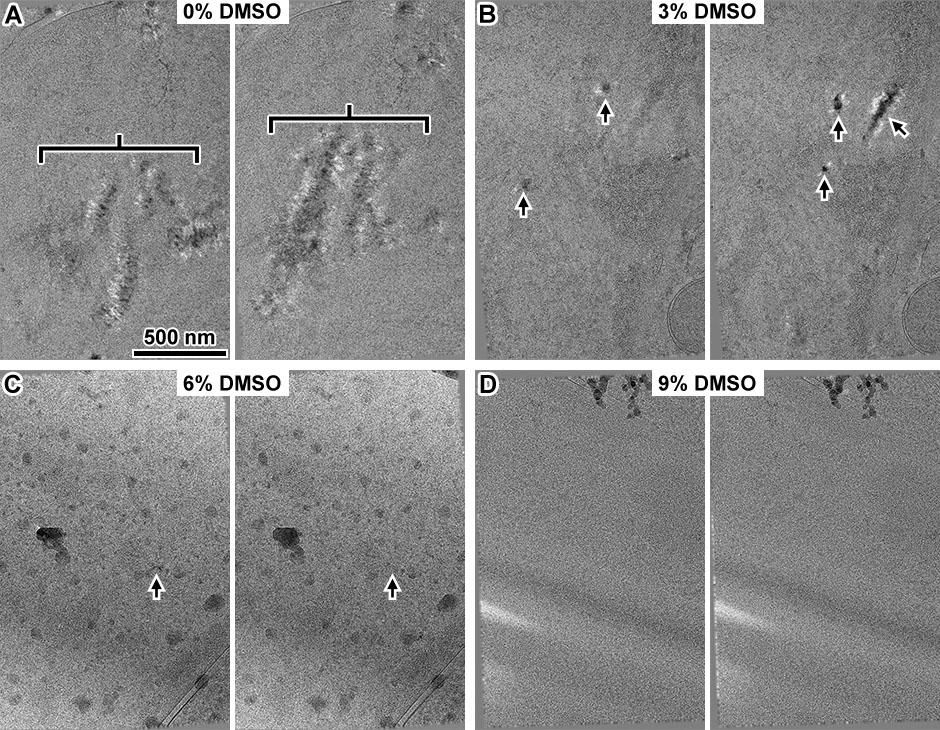

**Figure S4. Test of DMSO as a cryoprotectant for cryolamellae.**

Example G1-arrested RPE-1 cells, grown on EM grids, were briefly dipped in PBS+Ca+Mg supplemented with (A) 0%, (B) 3%, (C) 6%, or (D) 9% DMSO. The cells were then plunge-frozen, cryo-FIB milled, and imaged as a tilt series. To ensure that all four freezing conditions were tested on samples of similar thickness, the tilt series were collected at the nuclear periphery. Distinct nuclear double membranes (nuclear envelope) were observed for all tilt series, indicating that the sampled heights were indeed at the nuclear periphery. Note that the double membranes are not clear in panels B and D but are visible in the reconstructed cryotomograms. For each image pair, the tilt angle differed by 2°. The ice-crystal diffraction-contrast features are indicated by the brackets in panel A and short arrows in panel B. In panel C, the dense ~100- to 200-nm rounded features are ice-crystal contaminants on the lamella’s surface. In panel D, the bright and dark streaks oriented 10 to 4 o’clock are from uneven milling.

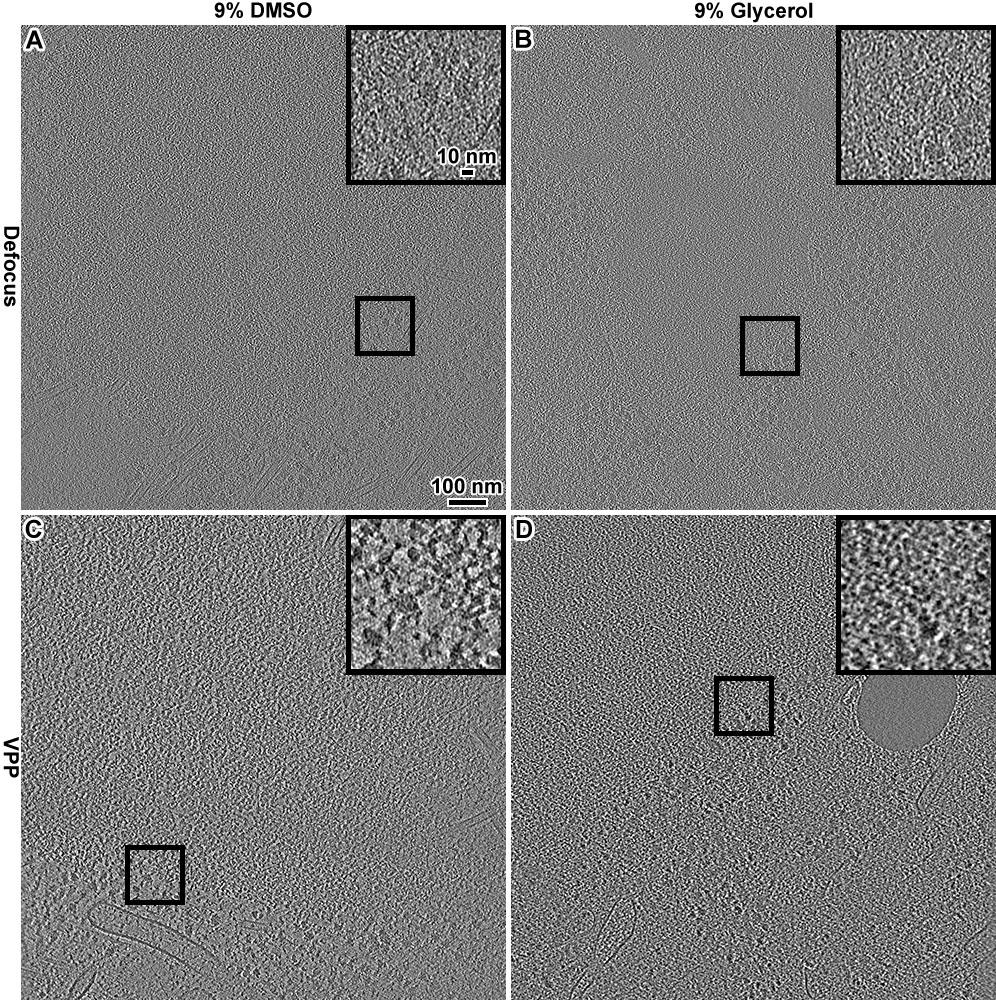

**Figure S5. Tests of intracellular cryo-ET contrast in cryoprotectants.**

Cryotomographic slices (12 nm) of metaphase-arrested RPE-1 cells frozen in (A) 9% DMSO and imaged by defocus phase contrast (Defocus), (B) 9% Glycerol and imaged by defocus phase contrast, (C) 9% DMSO and imaged with a Volta phase plate (VPP), (D) 9% Glycerol and imaged with a VPP. The fitted defocus was −5 µm in panels A and B and the nominal defocus was −0.5 µm in panels C and D. Insets show 3-fold enlargements of the corresponding boxed areas. A region that has both ribosomes and chromatin was chosen for each inset.

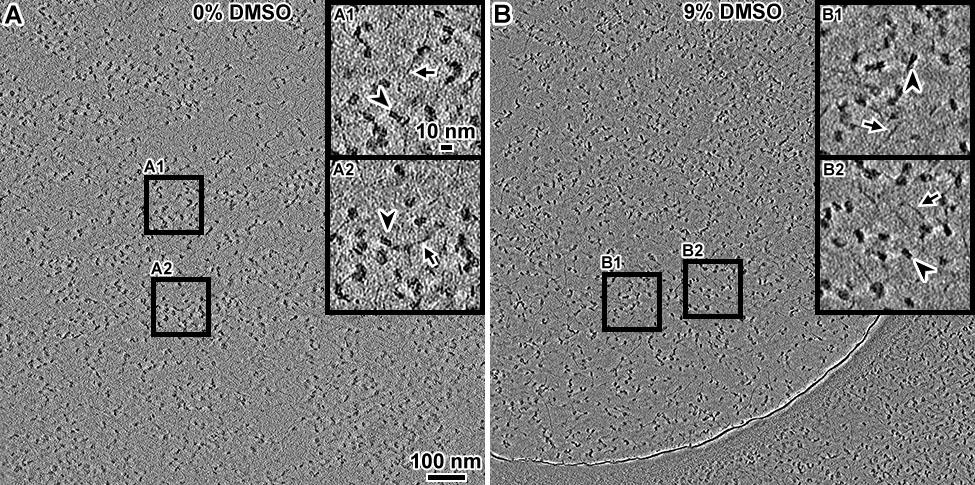

**Figure S6. Control cryo-ET of oligonucleosomes in DMSO cryoprotectant.**

Volta cryotomographic slices (10 nm) of HeLa oligonucleosomes in (A) storage buffer and (B) storage buffer plus 9% v/v DMSO. The granular densities are the nucleosomes. The large arc-shaped feature in the lower portion of panel B is the edge of the holey-carbon support film. The insets show 3-fold enlargements of boxed regions. Stretches of naked DNA (short arrows) and nucleosome double-gyre motifs (arrowheads) are indicated.

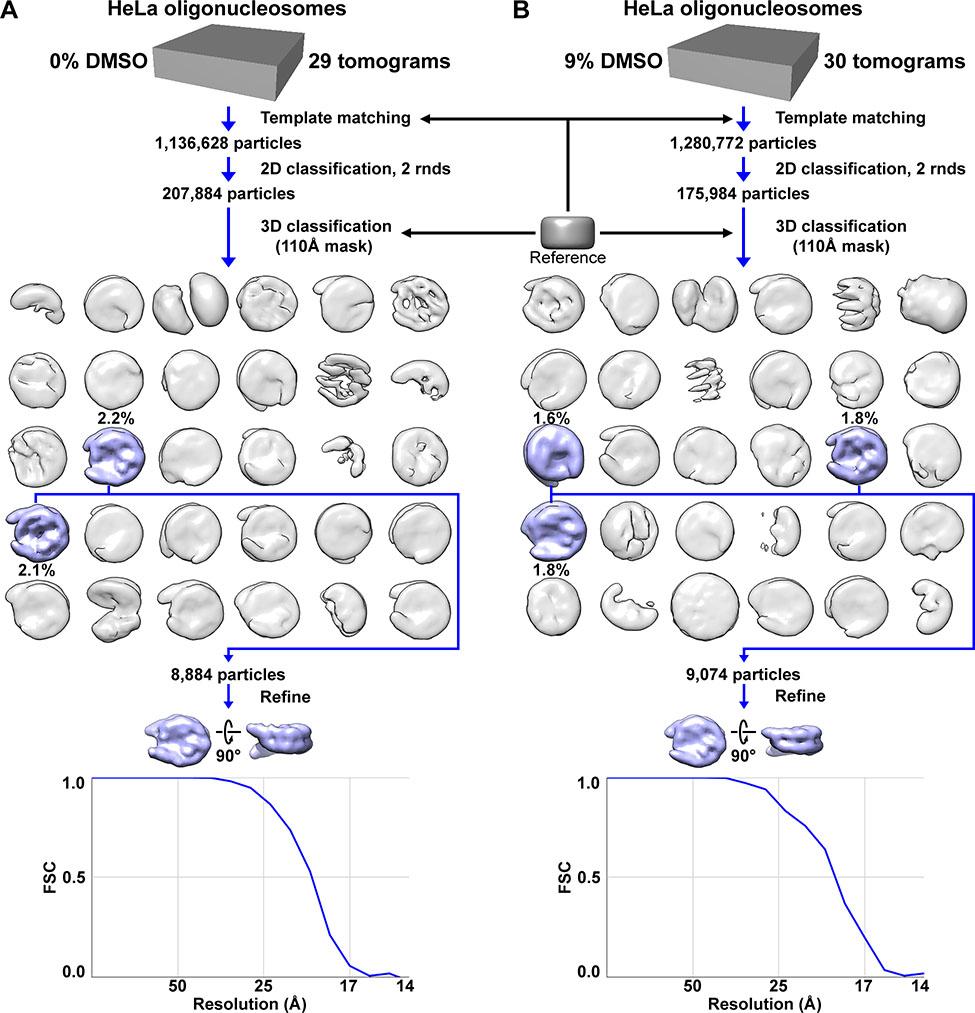

**Figure S7. Classification of HeLa oligonucleosomes in 0% and 9% DMSO.**

Classification and subtomogram analysis of oligonucleosomes in (A) 0% DMSO control and (B) 9% DMSO. Nucleosome-like particles were template matched using a 10-nm-wide, 6-nm-thick smooth cylindrical reference. Then two sequential rounds of 2-D classification were performed. One round of 3-D classification using k = 30 classes was done. The canonical nucleosomes with the higher-resolution features (those that didn’t have smooth surfaces) were combined for 3-D refinement. Based on the Fourier shell correlation (FSC) = 0.5 cutoff criterion, the resolution of the 0% and 9%-DMSO-treated oligonucleosomes class averages were 19 Å and 19.5 Å, respectively.

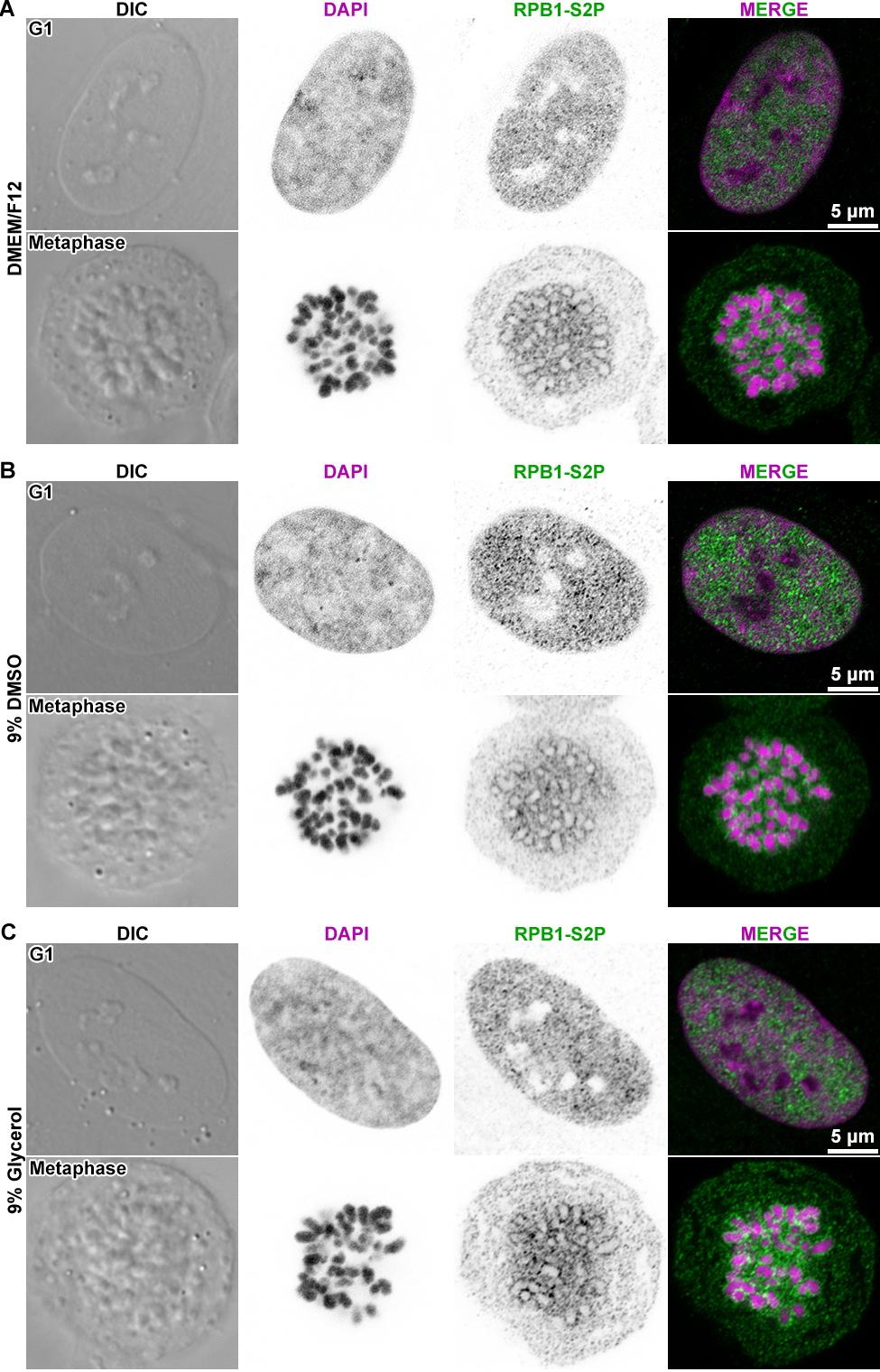

**Figure S8. Metaphase chromosomes are depleted of RNA polymerase II.**

Differential interference contrast (DIC) images of representative G1 and metaphase cells that were stained to detect DNA (stained with DAPI) and immunofluorescent detection of elongating RNAPII phosphorylated at serine 2 of the RPB1 subunit’s C-terminal tail heptad repeats (RPB1-S2P). (A) G1 and metaphase cells were incubated in DMEM/F12 medium for 1 minute before fixation. (B) G1 and metaphase cells were incubated in DMEM/F12 medium containing 9% DMSO for 1 minute before fixation. (C) G1 and metaphase cells were incubated in DMEM/F12 medium containing 9% glycerol for 1 minute before fixation. In the middle two columns, the DAPI and immunofluorescence signals are shown in inverted contrast for clarity.

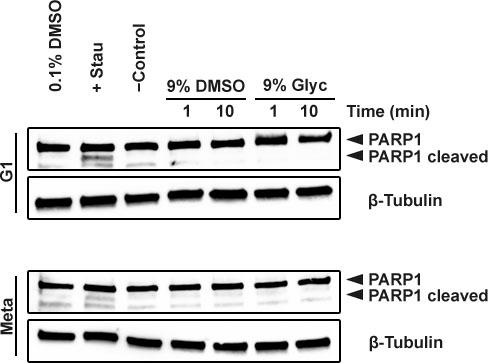

**Figure S9. Cryoprotection treatments do not induce apoptosis markers.**

Rows 1+2 and 3-4: Immunoblots of RPE-1 G1 and metaphase (Meta) cells, respectively. Rows 1 and 3 were probed for PARP1 while rows 2 and 4 were probed for beta-tubulin (loading control). The cleaved PARP1 band is found in apoptotic cells. Lanes 1 and 2: Apoptosis positive control in which RPE1 cells were treated with either carrier (0.1% DMSO) or 1 μM staurosporine (Stau) in 0.1% DMSO for 6 hours. Lane 3: untreated RPE-1 cells (negative control). Lanes 4 and 5: RPE-1 cells treated for 1 or 10 minutes with 9% DMSO. Lanes 6 and 7: RPE-1 cells treated for 1 or 10 minutes with 9% glycerol (Glyc). The uncropped blots are shown in Figure S36, C and D.

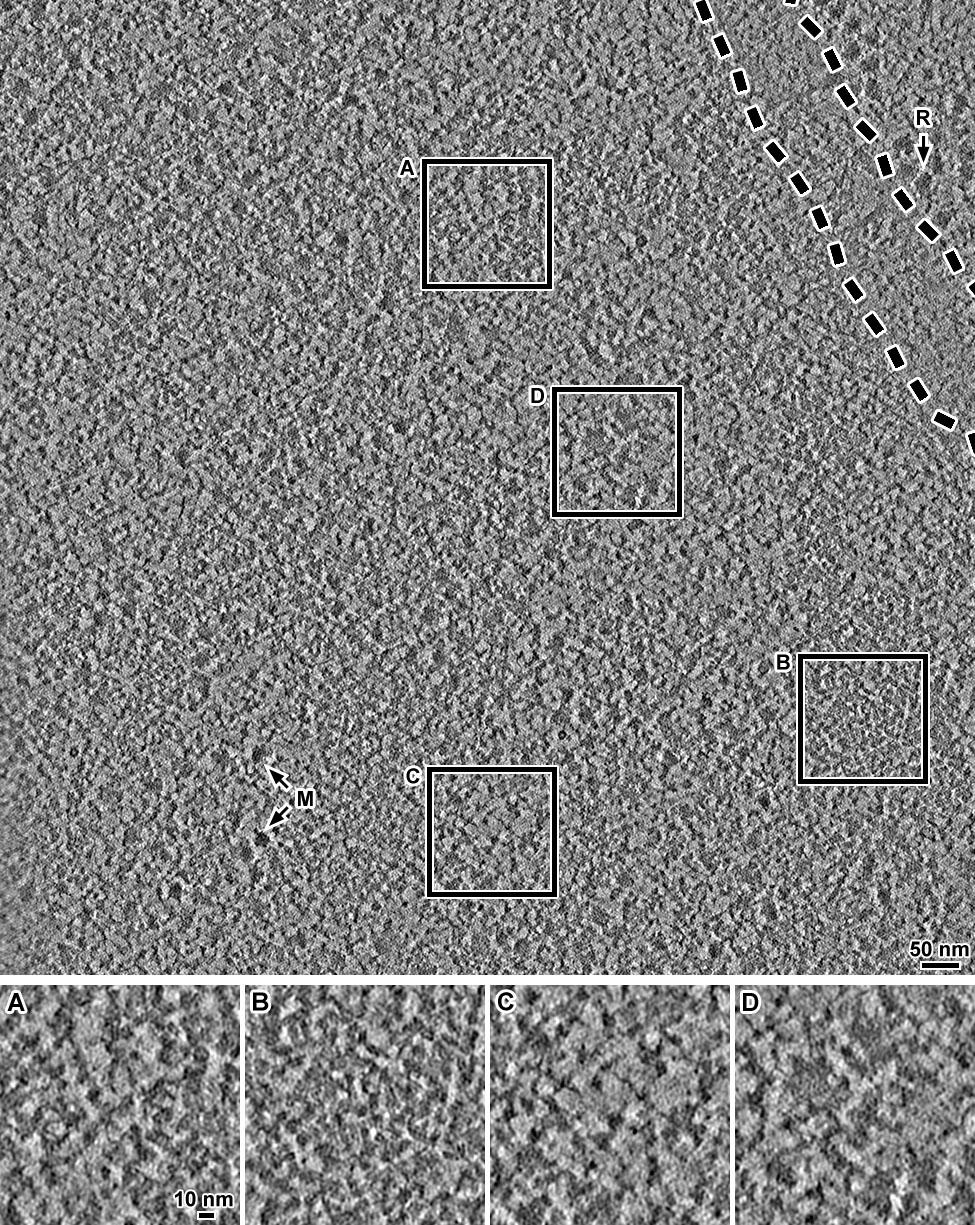

**Figure S10. Additional example Volta cryotomographic slice of a G1 cell.**

Cryotomographic slice (20 nm) of nuclear densities centered near the nuclear envelope of an RPE-1 cell. Rendered with low JPEG compression. Insets show 2-fold enlargements of (A and B) portions of chromatin domains, (C) region with fewer macromolecular complexes, and (D) a region with many megacomplexes. Cytological features are highlighted: megacomplex (M); ribosome (R). The region between the two dashed lines is the lumen of the nuclear envelope, which is oriented obliquely to the milling direction. As such, the nuclear envelope membranes are not visible.

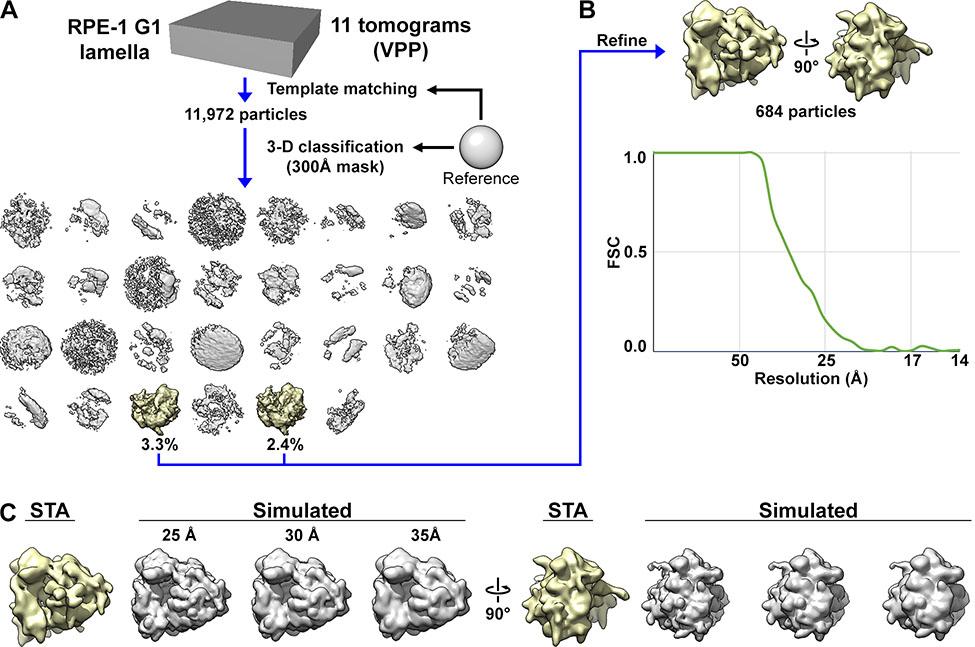

**Figure S11. Controls subtomogram analysis of ribosomes *in situ*.**

(A) Template matching was done only on the cytoplasmic regions, using a spherical reference. The template-matching hits were then subjected to direct 3-D classification (also using a spherical reference), revealing numerous non-ribosome class averages (gray) and two 80S ribosome class averages (yellow). (B) The particles of these two class averages were combined and refined to ~33 Å resolution, based on the FSC = 0.5 criterion. (C) The (yellow) refined *in situ* 80S ribosome density map reproduced from panel B, alongside (gray) density maps of purified human ribosomes (4UG0) (Khatter *et al*, 2015) simulated at different resolutions.

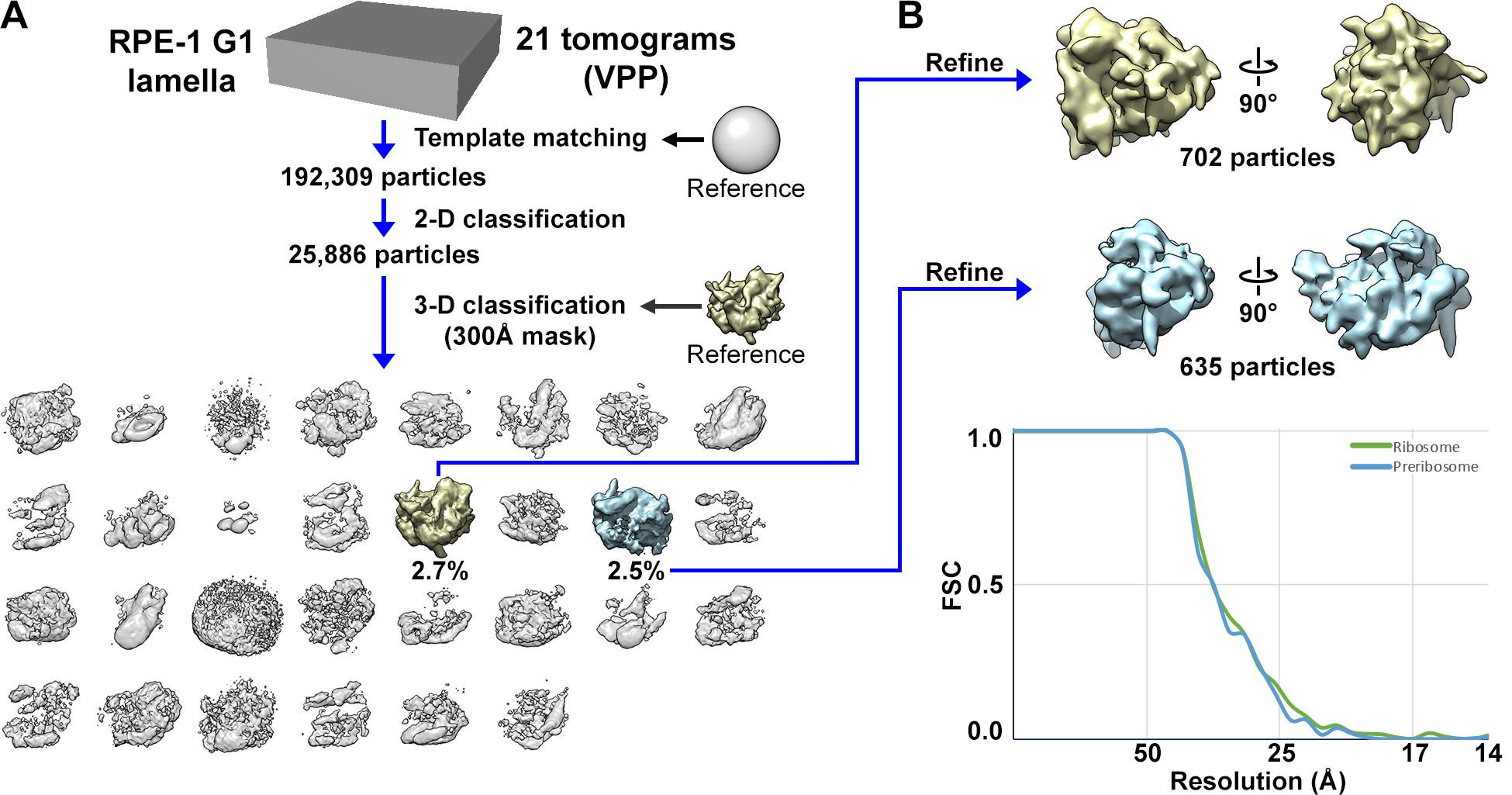

**Figure S12. Subtomogram analysis of preribosomes.**

(A) Template matching for preribosomes was performed on all cryotomograms containing nuclear regions, using a spherical reference. The candidate hits were then subjected to 2-D classification; classes that contain subtomograms that do not correspond to large complexes were removed. The remaining subtomograms were then subjected to 3-D classification, using the ribosome refined class average shown in Figure S11B as the reference, but low-pass filtered to 60 Å resolution. (B) The preribosome (blue) class average was refined to ~32 Å resolution, based on the FSC = 0.5 criterion. The 80S ribosome average (yellow) is also shown for comparison purposes. The preribosome resembles the 60S subunit of the mature ribosome and is oriented to better show their similarities.

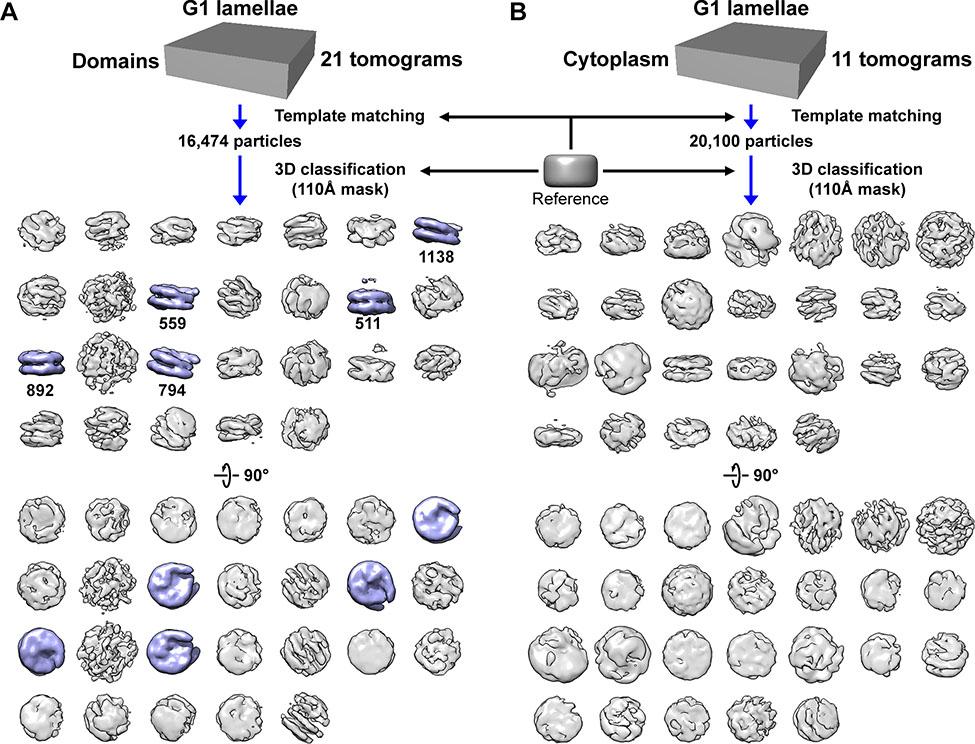

**Figure S13. Controls for nucleosome 3-D classification *in situ*.**

(A) Subtomogram analysis of G1 chromatin domains. (B) Subtomogram analysis of G1 cytoplasm. Template matching was done with a featureless cylinder reference and a 110 Å spherical mask. The grid spacing for both template matching experiments was 21 nm. The same reference and mask were used for both datasets. Canonical nucleosome class averages are shaded in blue while ambiguous class averages are shaded in gray. The ambiguous densities (gray) are not canonical nucleosomes; they are abundant because the template-matching process uses a featureless cylinder reference and a low cross-correlation cutoff. As a result, large numbers of false positives are rejected in the classification analysis. Classification was done using 30 classes for each experiment. Some classes had too few particles and were therefore not shown.

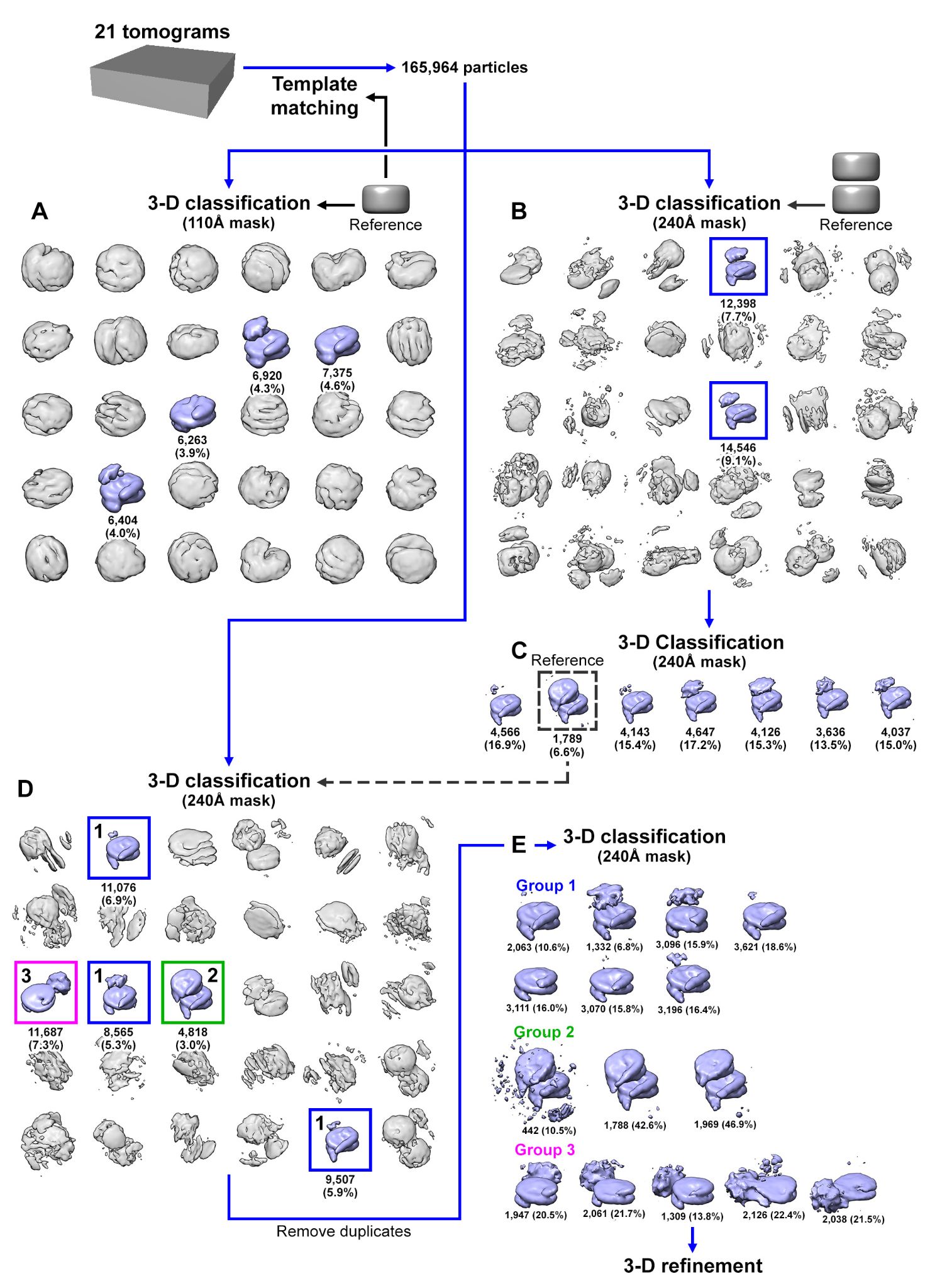

**Figure S14. Classification flowchart of G1 chromatin domains.**

(A) G1 candidate nucleosome subtomograms that were template matched with a cylindrical reference were directly classified in 3-D, using a cylindrical reference and a 110 Å spherical mask. Note that there are more template-matching hits in this set of experiments than that of Figure S13 because a smaller grid spacing was used here. The ambiguous densities (gray) are not canonical nucleosomes; they are abundant because the template matching process uses a featureless cylinder reference and a low cross-correlation cutoff. As a result, large numbers of false positives are rejected in the classification analysis. (B) In parallel, the same set of subtomograms were directly classified in 3-D using a stacked cylinder reference and a larger spherical mask. (C) A second round of classification using a nucleosome class average from panel B as the reference and the larger mask yielded mononucleosomes with extra densities at their face plus an unambiguous dinucleosome class. (D) Direct 3-D classification was done on the original set of subtomograms using a larger mask and the stacked dinucleosome class average from panel C as the reference. Three groups of class averages were obtained, corresponding to (1) mononucleosome, (2) stacked dinucleosome, and (3) mononucleosome with a gyre-proximal density. (E) These three groups of class averages were subjected to a third round of classification using either a mononucleosome, stacked dinucleosome or mononucleosome with a gyre-proximal density from panel D as the reference and large spherical mask.

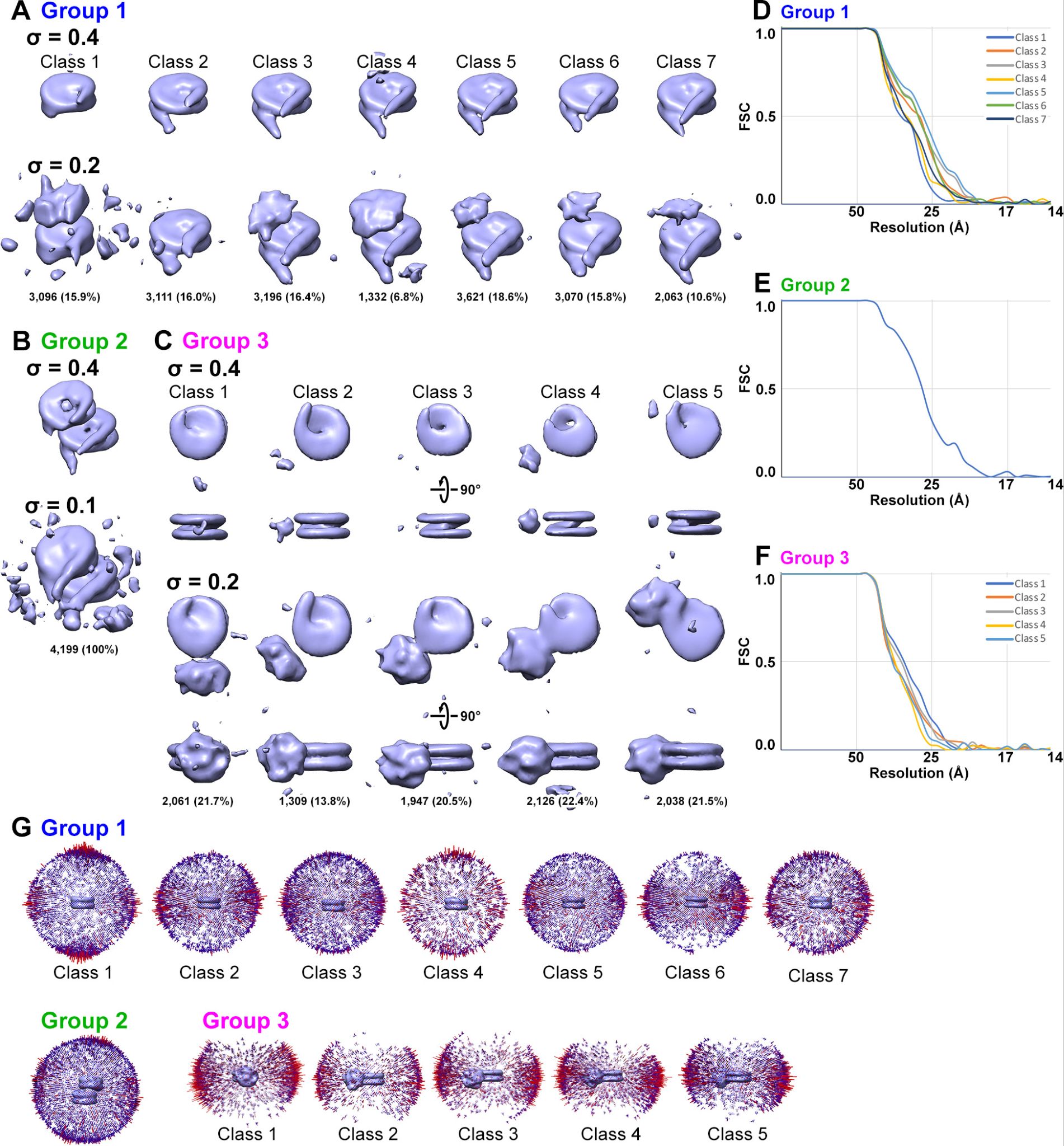

**Figure S15. Refinement of G1 mononucleosomes and dinucleosomes.**

Refined class averages for (A) mononucleosomes, (B) stacked dinucleosomes, and (C) mononucleosomes with a gyre-proximal density. Two contour levels are shown for each class average. (D, E, F) FSC plots and (G) angular distribution of the refined density maps. For panels A – C, the hide dust feature in UCSF Chimera was disabled.

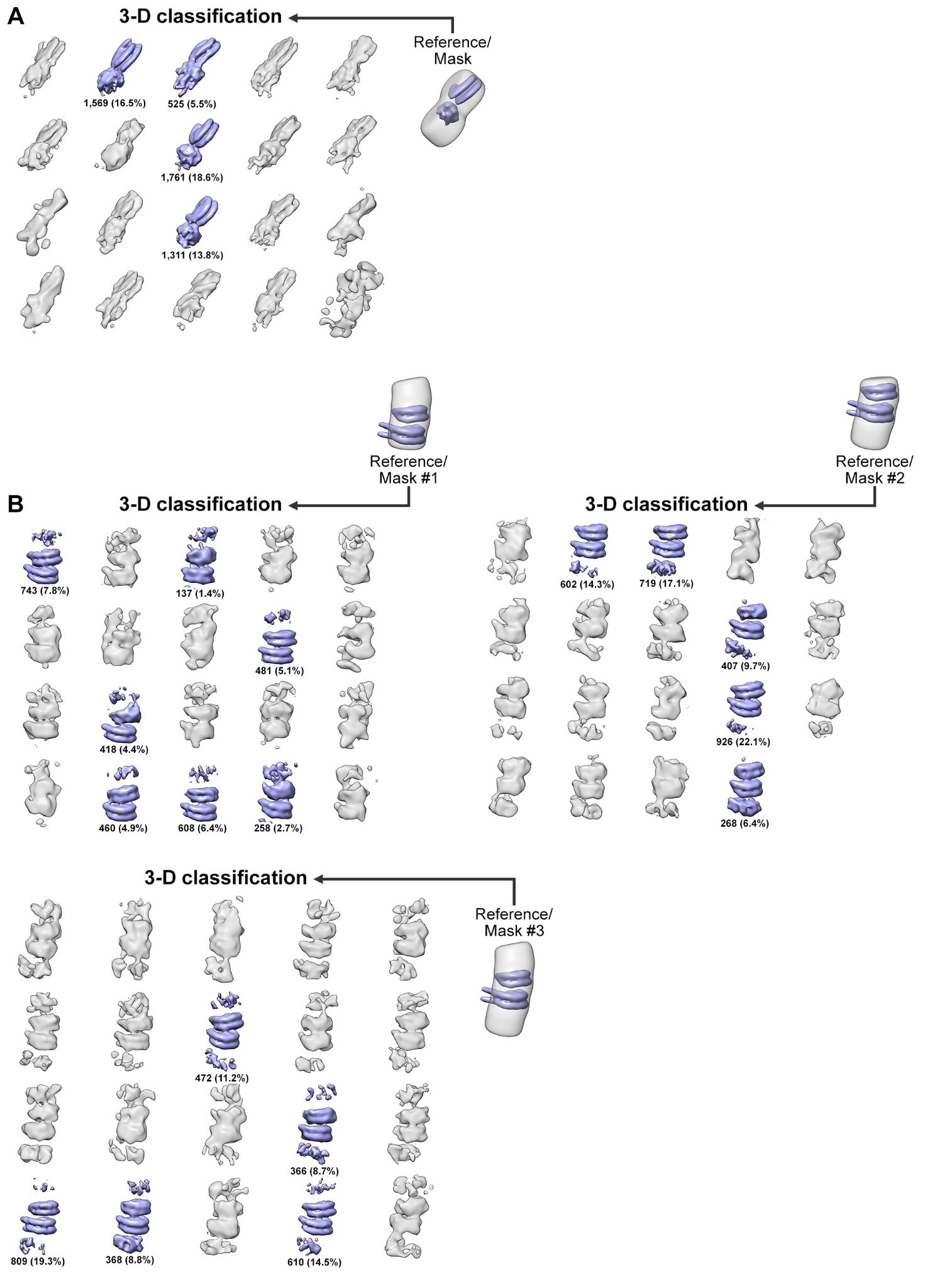

**Figure S16. Subtomogram analysis of alternative ordered nucleosome packing motifs in G1 cells.**

Nucleosome particles from Groups 2 and 3 (Figures S15 B and C, respectively) were subjected to an additional round of 3-D classification, using custom masks that enclose volumes where an additional complex may reside. Since the particles were already aligned from the previous refinement step, a restricted angular search range was imposed for these runs. The “reference/mask” models in the figure depicts the location of the volume masked-in (gray), relative to the reference (blue) used for each classification run. The masks used for these 3-D classification runs were optimized for (A) side-by-side nucleosomes and (B) ordered trinucleosomes and tetranucleosomes. Class averages that contain at least one ordered nucleosome are shaded blue.

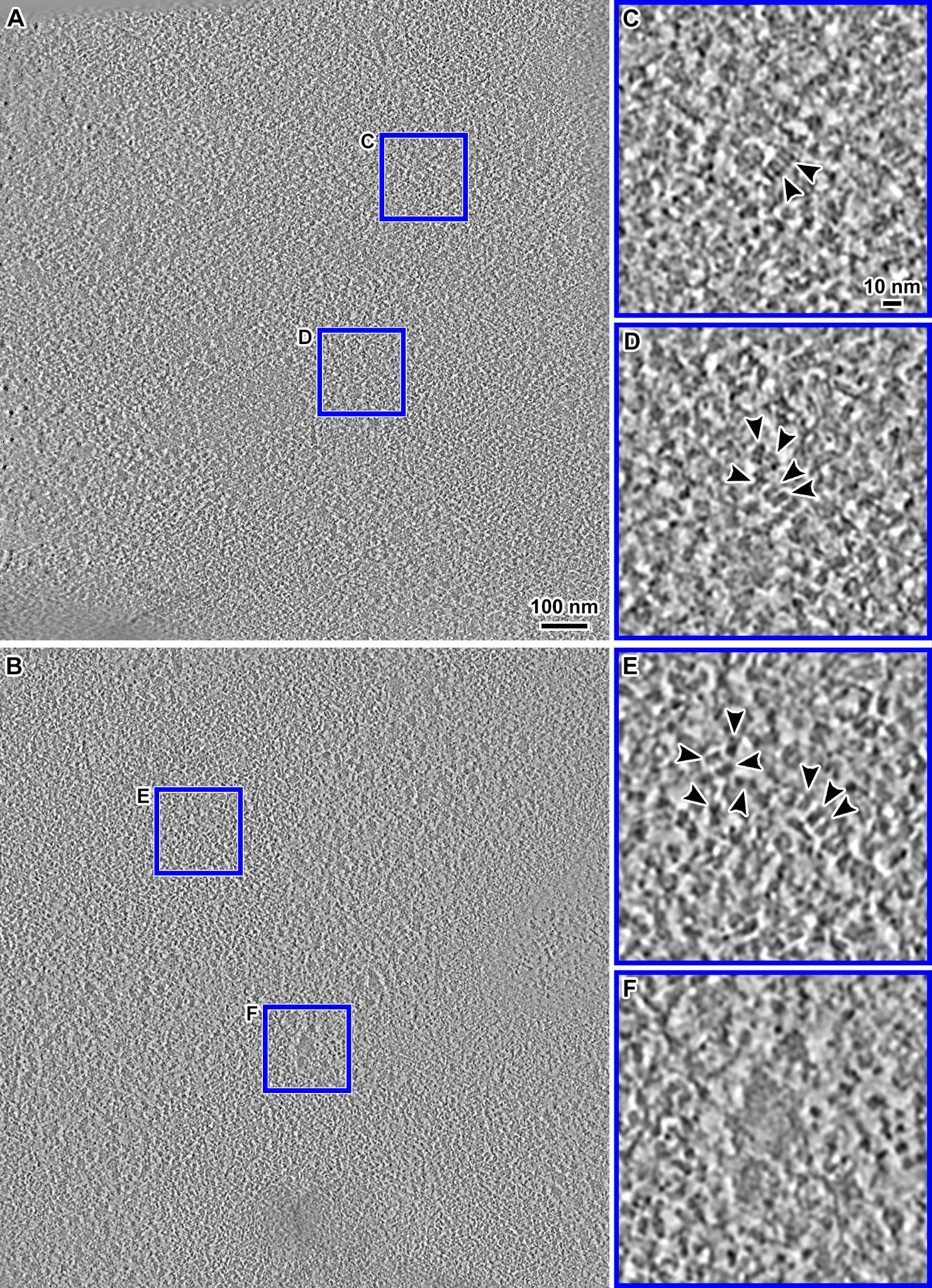

**Figure S17. Cryo-ET of G1 chromatin *in situ* of cells with glycerol cryoprotection.**

(A & B) Cryotomographic slices (10 nm) of the nuclear region of G1 RPE-1 cells that were plunge frozen in the presence of 9% glycerol for cryoprotection. (C & D) Enlargements (4-fold) of the two boxed regions in panel A, showing chromatin domains. (E) Enlargement (4-fold) of another chromatin domain in Panel B. (F) Enlargement (4-fold) of a “dense irregular body” in panel B. Nucleosomes are indicated by black arrowheads in panels C, D and E. These images are from cryotomograms denoised by CryoCARE (Buchholz *et al*, 2019).

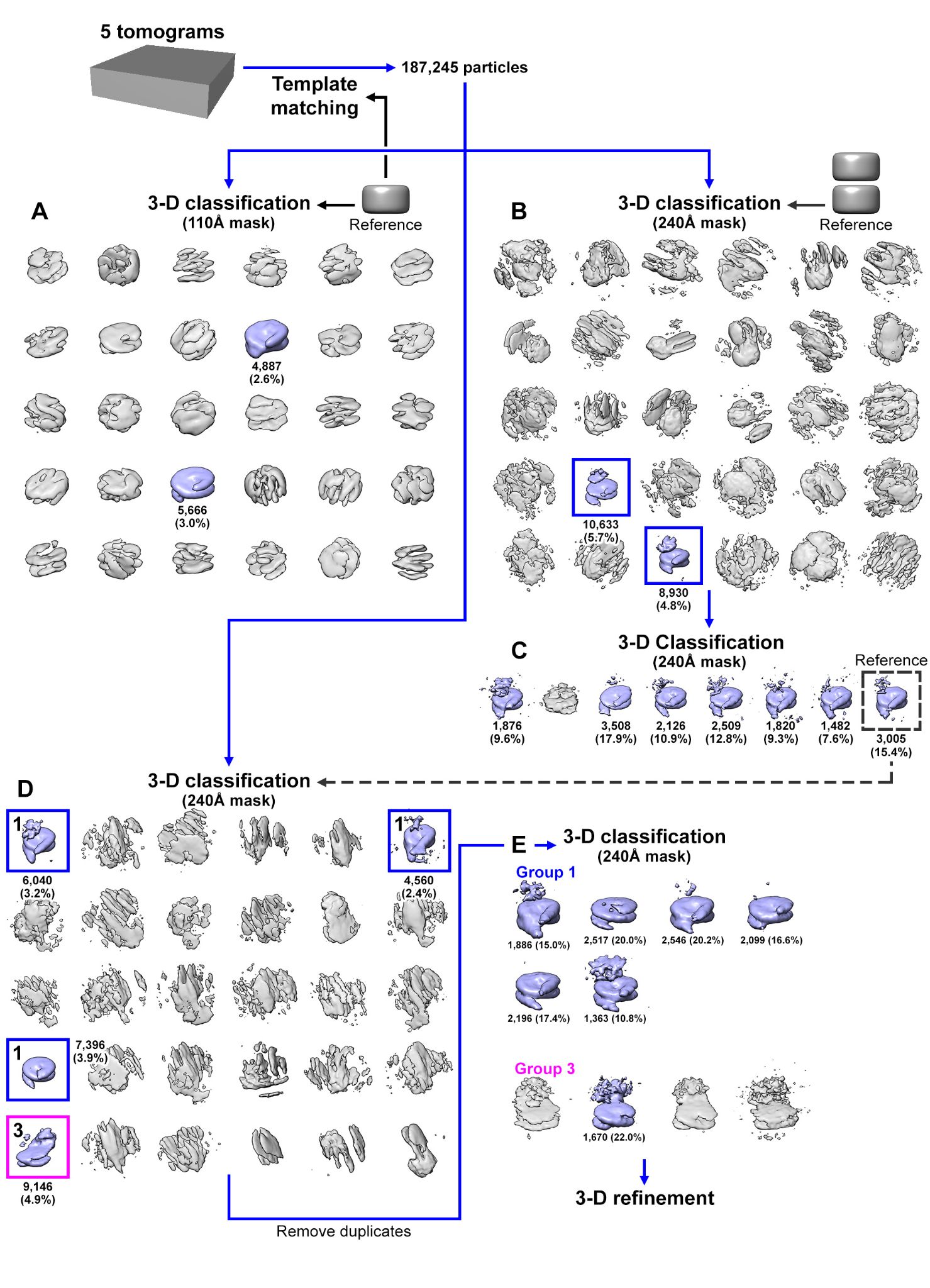

**Figure S18. Classification flowchart of G1 chromatin domains in 9% glycerol-cryoprotectant cells.**

The same classification workflow used in the analysis of the 9% DMSO-cryoprotected cell dataset (Figure S14) is used here. Briefly, tomograms collected from 9% glycerol-cryoprotected G1 cells were template matched using a featureless cylindrical reference. (A) The resultant candidate nucleosome subtomograms were then classified in 3-D, using the cylinder reference and a 110 Å spherical mask. (B) In parallel, the same set of candidate subtomograms were also classified using a stacked cylinder reference and a larger 240 Å spherical mask. (C) Nucleosome class averages detected in panel B were then subjected to a second round of classification, using one of the class averages as the reference. Unlike in the tomograms of 9% DMSO-cryoprotected cells (Figure S14C), the stacked dinucleosome class average was not detected here. (D) The original set of the candidate subtomograms was classified again, this time using a mononucleosome class average from panel C as the reference. Two groups of mononucleosome class averages were obtained, corresponding to (Group 1, boxed blue) mononucleosomes and (Group 3, boxed magenta) mononucleosomes with a proximal-gyre density. (E) The two groups were then subjected to a third round of classification separately, using either a mononucleosome or a mononucleosome with proximal-gyre density class average from panel D as the reference. The nucleosome class averages in panel E were then individually subjected to refinement (see Figure S19). Nucleosome class averages are colored blue; ambiguous class averages are colored gray.

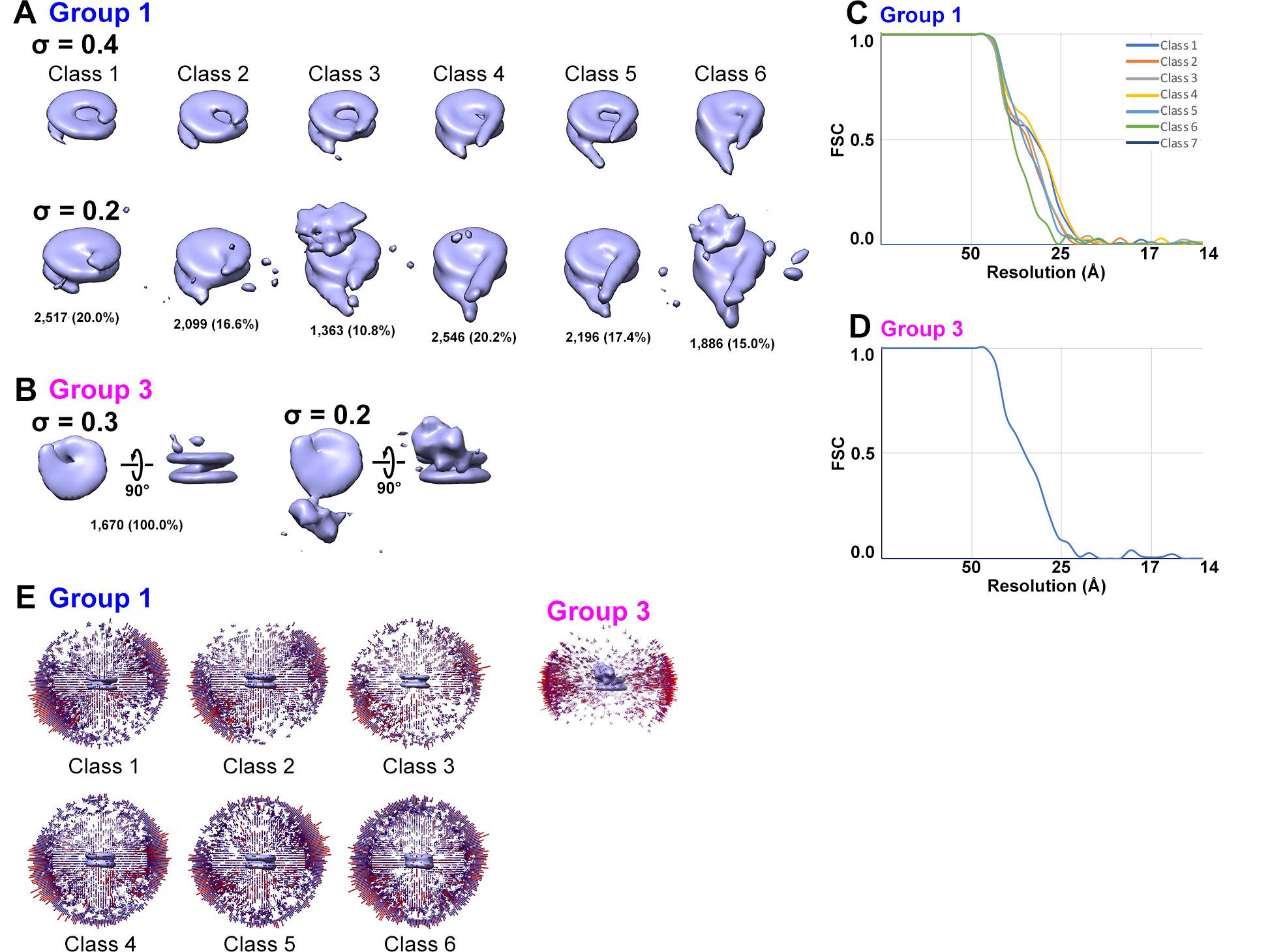

**Figure S19. Refinement of mononucleosomes from glycerol-cryoprotected G1 cells.**

Refined class averages for (A) mononucleosomes and (B) mononucleosome with a proximal-gyre density. Two contour levels are shown for each class average. (C & D) FSC plots and (E) angular distributions of the refined class averages in panels A and B.

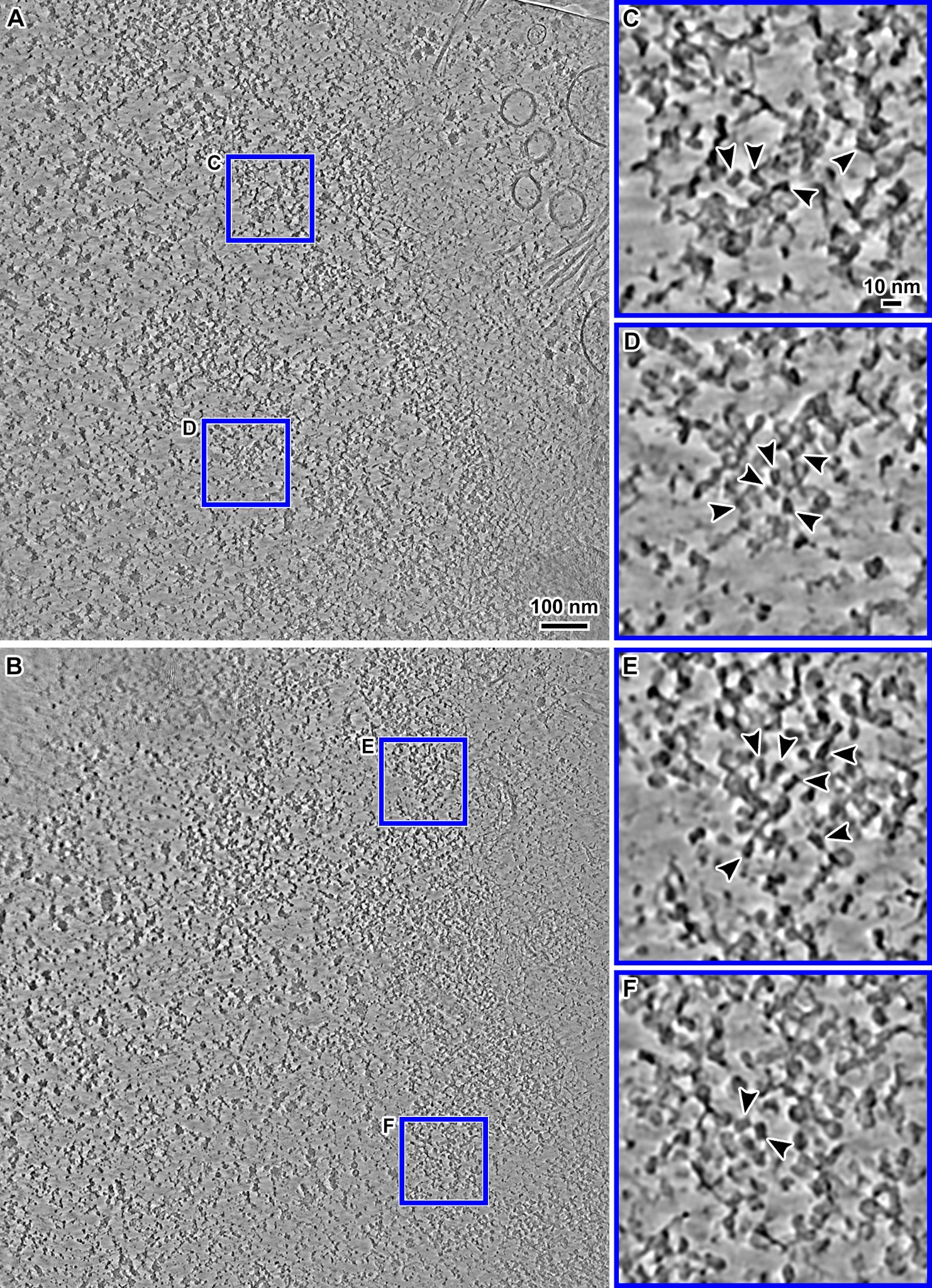

**Figure S20. Cryo-ET of G1 chromatin *in situ* of cells without cryoprotection.**

(A & B) Cryotomographic slices (10 nm) of perinuclear regions of G1 RPE-1 cells that were plunge frozen without addition of a cryoprotectant. The nuclear envelopes are indicated by the dashed lines. (C & D) Enlargements (4-fold) of chromatin domains in Panel A. (E & F) Enlargements (4-fold) of chromatin domains in panel B. Nucleosomes in panels C – F are indicated by black arrowheads. These images are from cryotomograms denoised by CryoCARE (Buchholz *et al.*, 2019).

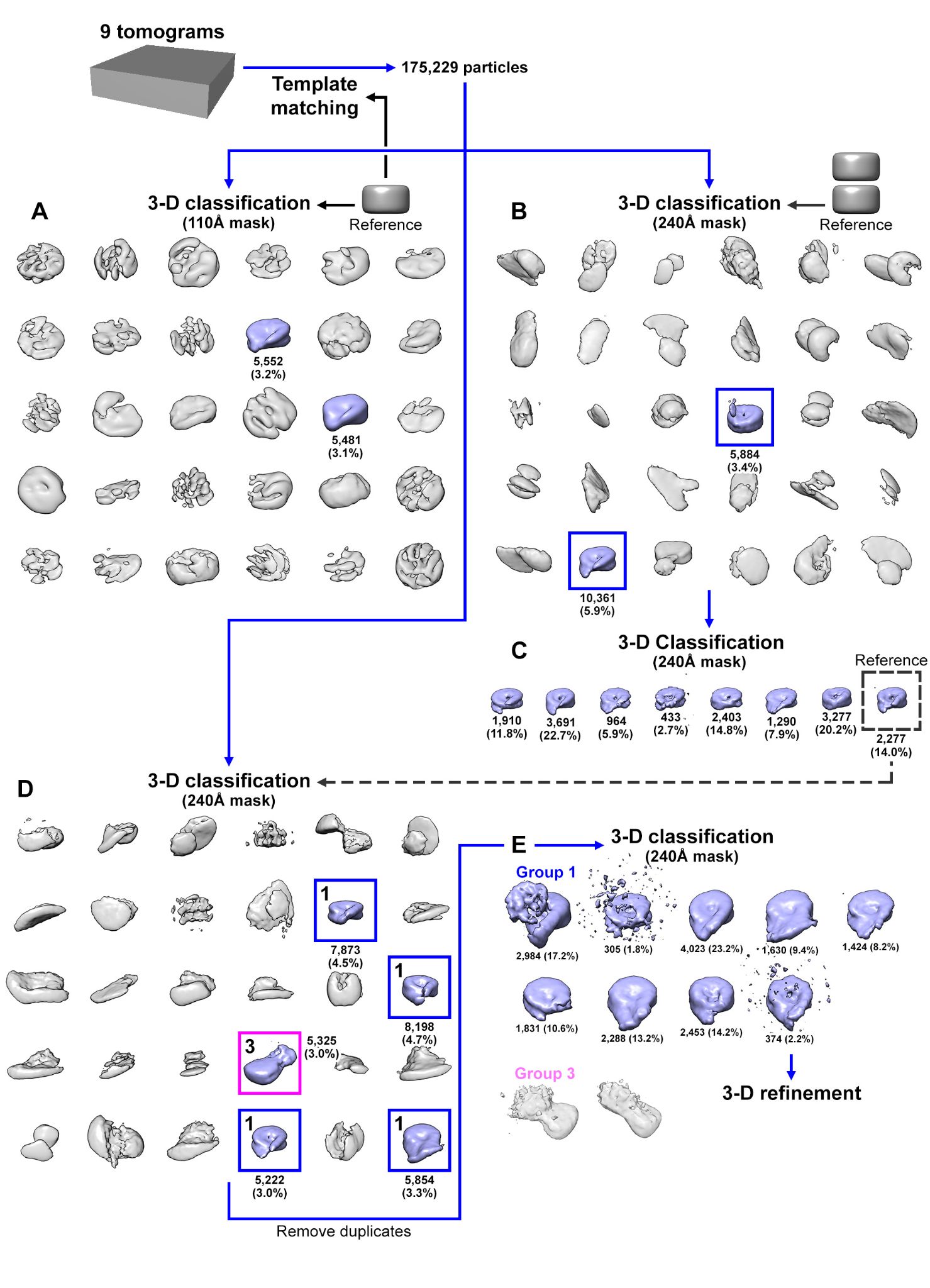

**Figure S21. Classification flowchart of G1 chromatin domains for cells without cryoprotection.**

The same classification workflow used in the 9% DMSO- (Figure S14) and 9% glycerol-cryoprotected (Figure S18) G1 datasets was used for the analysis here. Tomograms collected from G1 cells plunge frozen without cryoprotection were subjected to template matching using a featureless cylindrical reference. (A) Subsequently, the candidate nucleosomes from template matching were classified in 3-D using the cylindrical reference and a 110 Å spherical mask. (B) In parallel, the candidate nucleosomes were also classified using a stacked cylinder reference and a larger 240 Å spherical mask. (C) Nucleosome class averages detected in panel B were then subjected to a second round of 3-D classification, using one of the nucleosome class averages from Panel B as the reference. The stacked dinucleosome class average was not detected at this stage. (D) The original set of candidate nucleosomes was once again subjected to 3-D classification, this time using one of the nucleosome class averages from panel C (dashed box) as the reference. Several mononucleosome class averages (Group 1, boxed blue) and one class average resembling the mononucleosome with a proximal-gyre density (Group 3, boxed magenta) were detected here. (E) Nucleosomes in Group 1 and 3 were then separately subjected to another round of classification. The Group 1 mononucleosomes obtained in this round of classification were then subjected to refinement (Figure S22). For Group 3, the resultant class averages did not reveal a clear nucleosome density and thus were not analyzed further. Nucleosome class averages are colored blue; ambiguous class averages are colored gray.

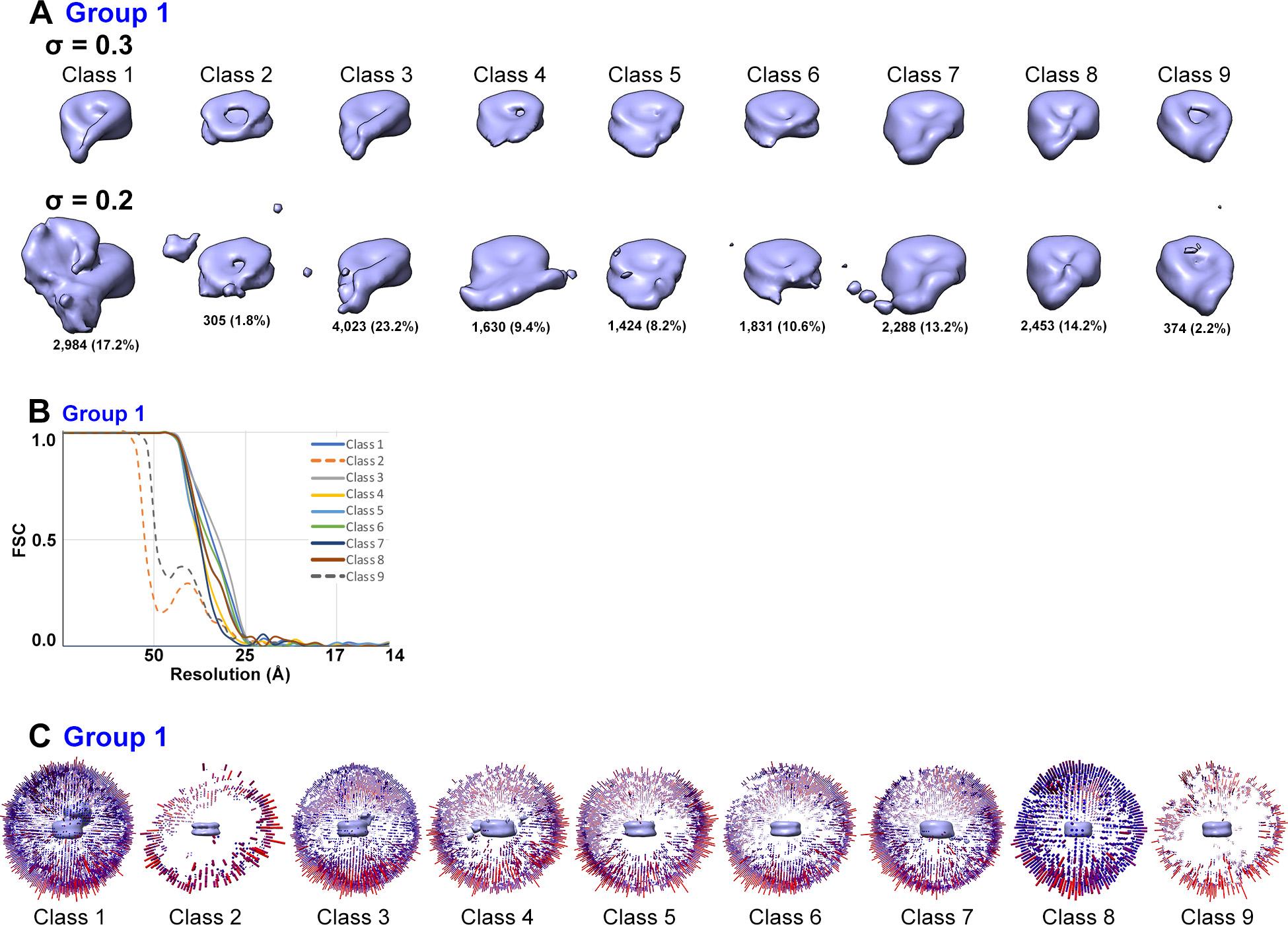

**Figure S22. Refinement of Group 1 nucleosome class averages from G1 cells without cryoprotection.**

(A) Refined mononucleosome class averages, rendered at two different threshold levels. The FSC plot and angular distributions of the class averages are shown in panels B and C, respectively.

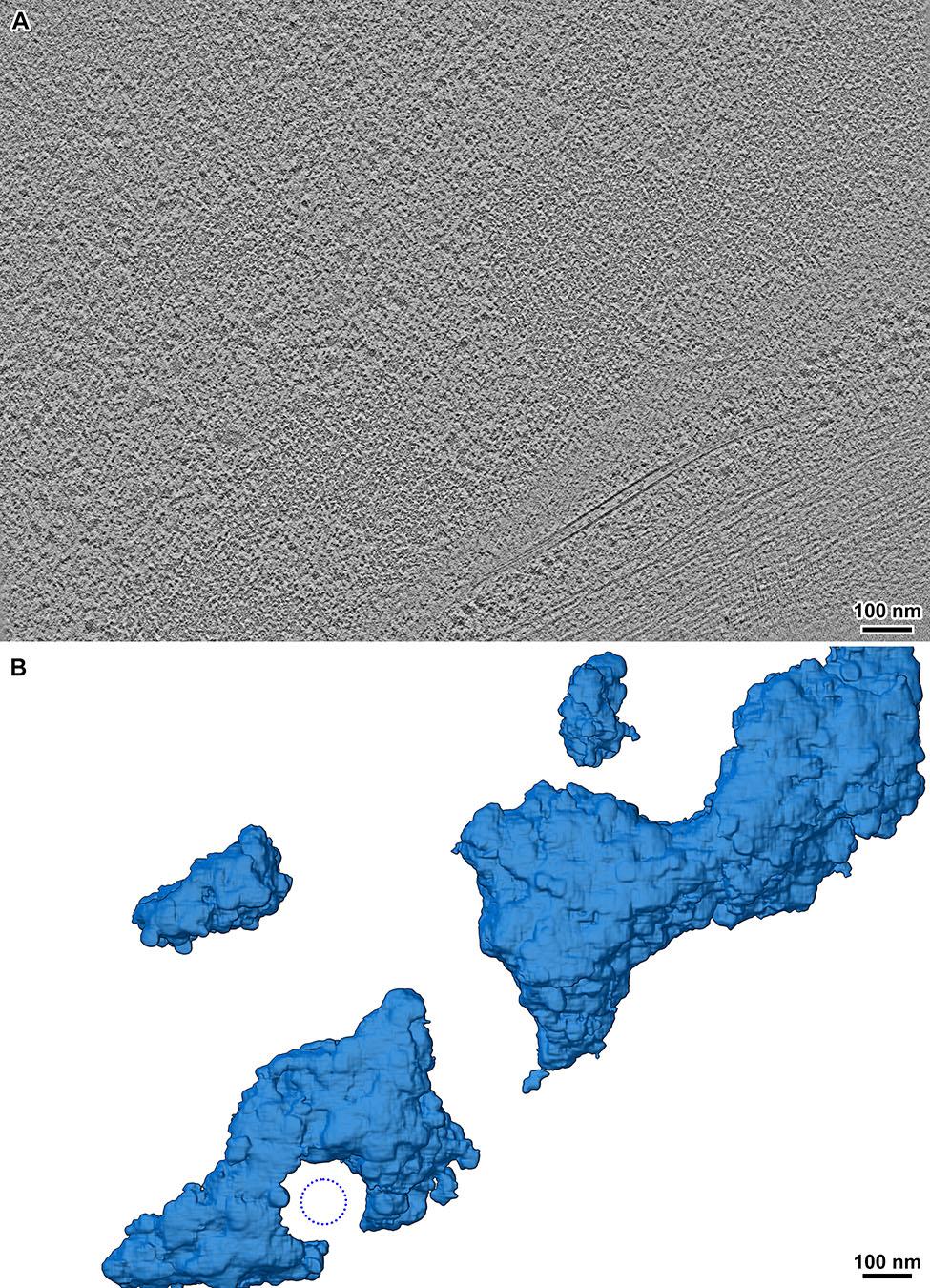

**Figure S23. G1 chromatin domains are irregular.**

(A) Cryotomographic slice a region near the nuclear envelope. (B) Convolutional neural network (CNN) based segmentation of the region in panel A. For clarity, the CNN segmentation rendering shows the entire ~90 nm thickness of the lamella whereas the cryotomographic slice shows the central 10 nm. Because of this difference, the CNN segmentation shows more chromatin than is visible in the cryotomographic slice. The dotted blue circle indicates the approximate position of the nuclear pore complex (not segmented).

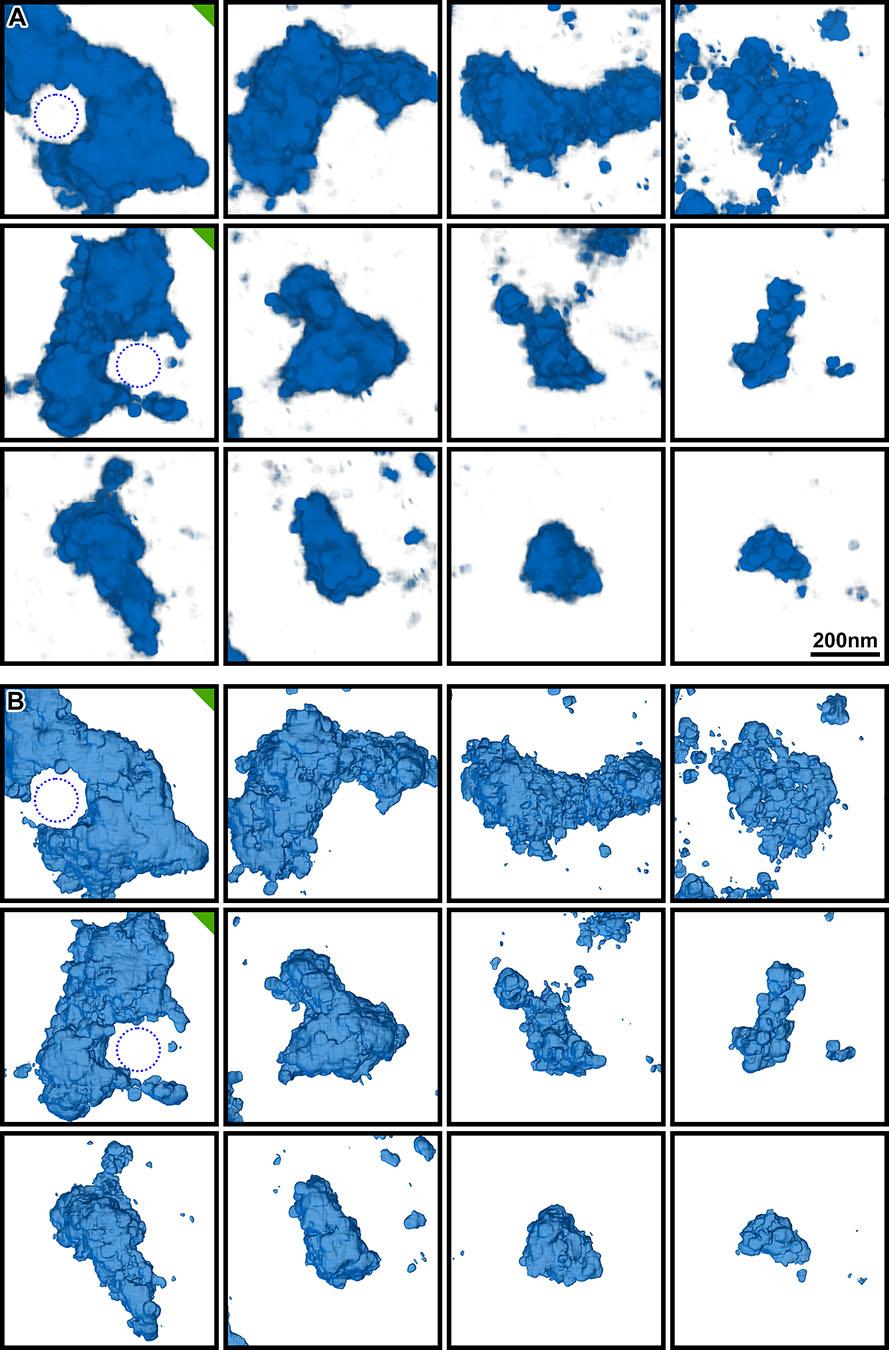

**Figure S24. Additional CNN annotations of chromatin domains in G1 nuclei.**

(A) Volume and (B) isosurface renderings of chromatin domains. Panels marked with a green triangle contain perinuclear chromatin domains. The dotted blue circles indicate the approximate positions of nuclear pore complexes (not segmented).

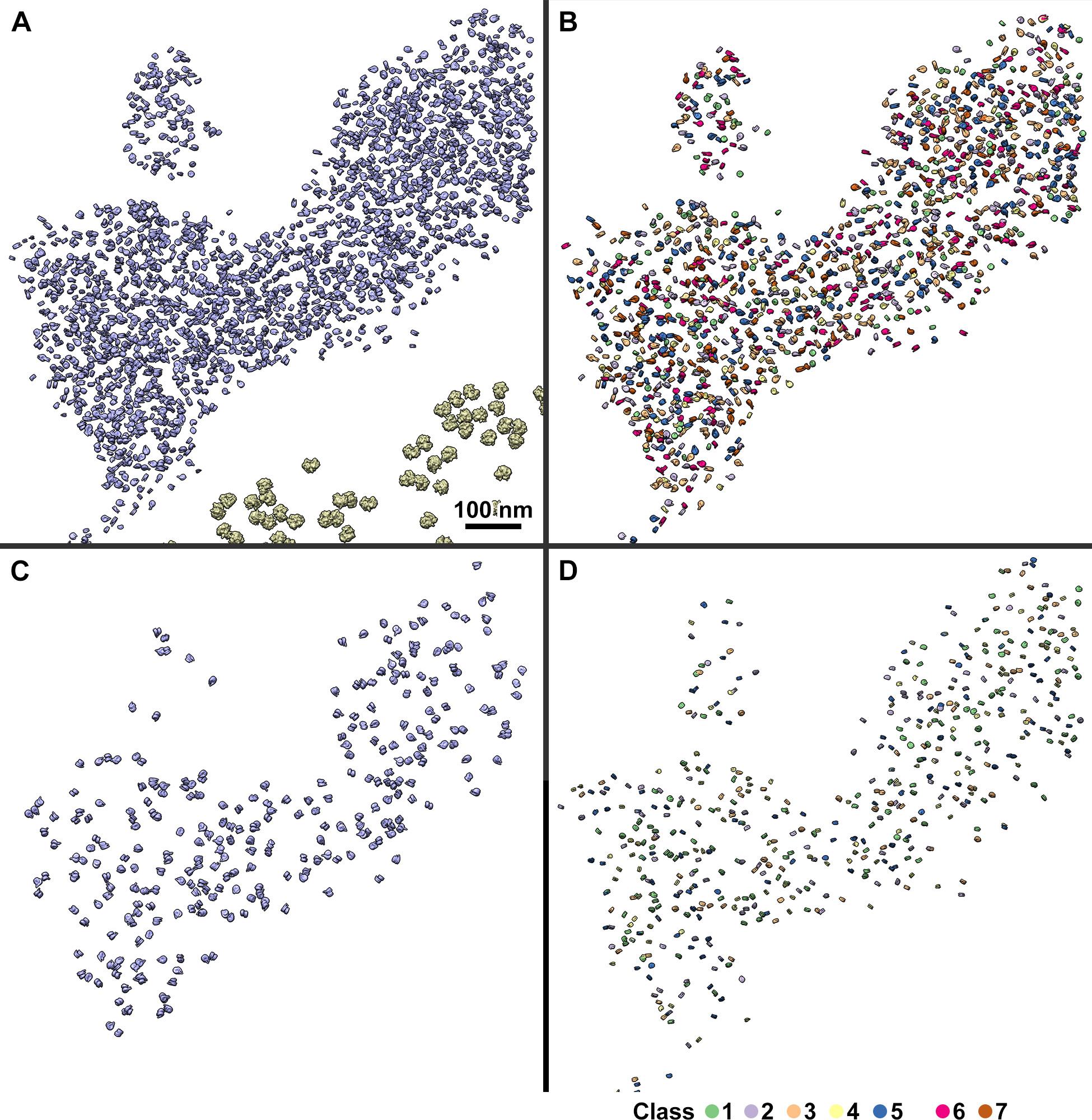

**Figure S25. Remapped models of G1 nucleosome groups.**

(A) Enlargement of a segment of Figure 3D showing all remapped nucleosomes (blue) in a G1 domain. Class averages of mononucleosomes (group 1), ordered stacked dinucleosomes (group 2) and mononucleosomes with gyre-proximal density (group 3) are remapped separately in panels B, C and D respectively. The color-coding of the nucleosome densities in panels B and D correspond to the class averages shown in Figure 3A and Figure 3C.

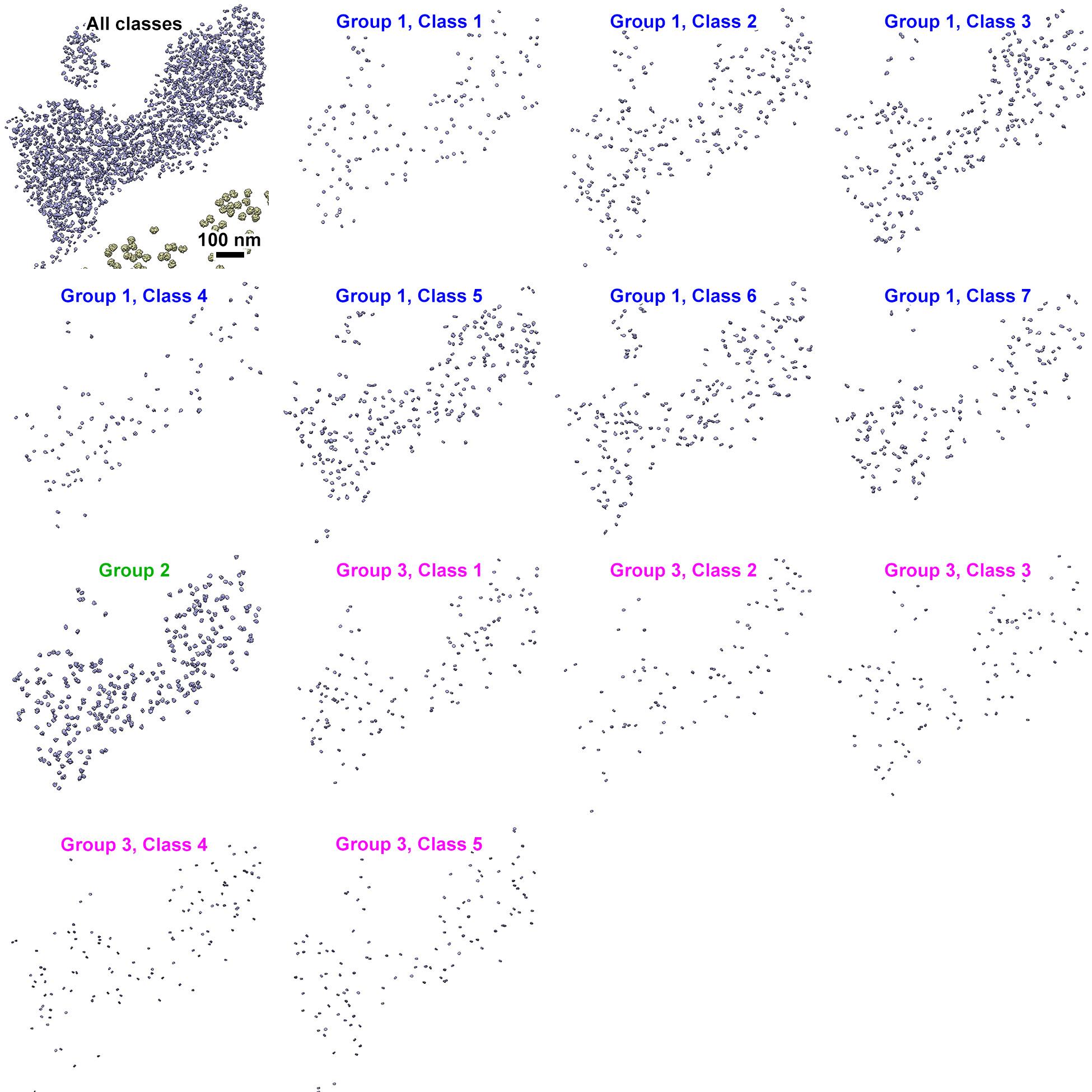

**Figure S26. Remapped models of G1 nucleosome individual classes.**

The upper left panel shows a section of the remapped model presented in Figure 3D. The remaining panels also show the same section of the remapped model, but with each of the nucleosome class averages from Figure 3A – C rendered separately.

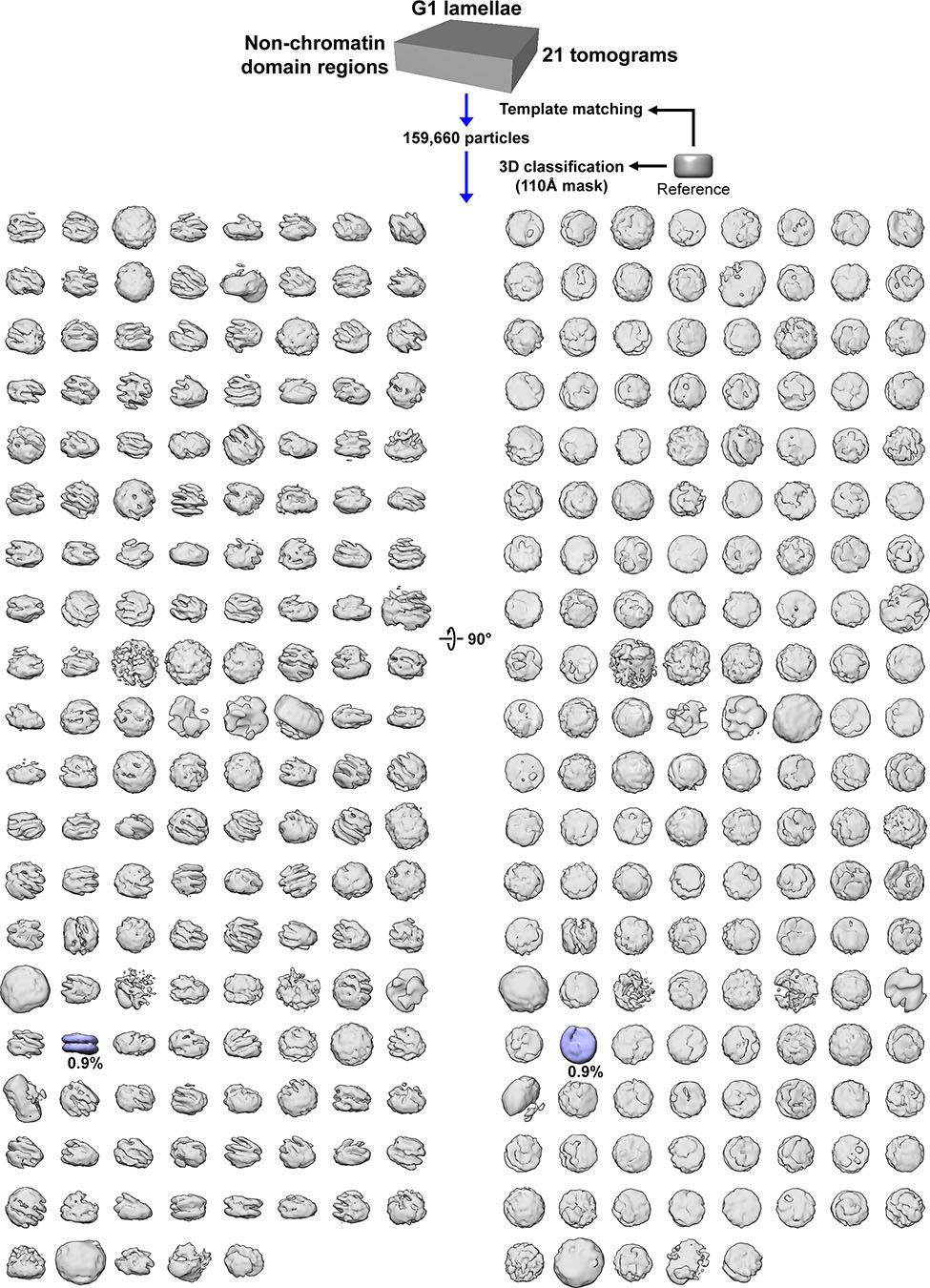

**Figure S27. Subtomogram analysis of the G1 nucleoplasm.**

The regions outside the chromatin domains were template matched with a featureless cylinder reference. The template-matching hits were then subjected to direct 3-D classification with 200 classes using the same featureless cylinder as the reference. Out of the 157 non-empty classes, the vast majority do not resemble canonical nucleosomes (gray). Only one canonical nucleosome class average was detected (blue), but the subtomograms from this class were mostly located at the periphery of chromatin domains.
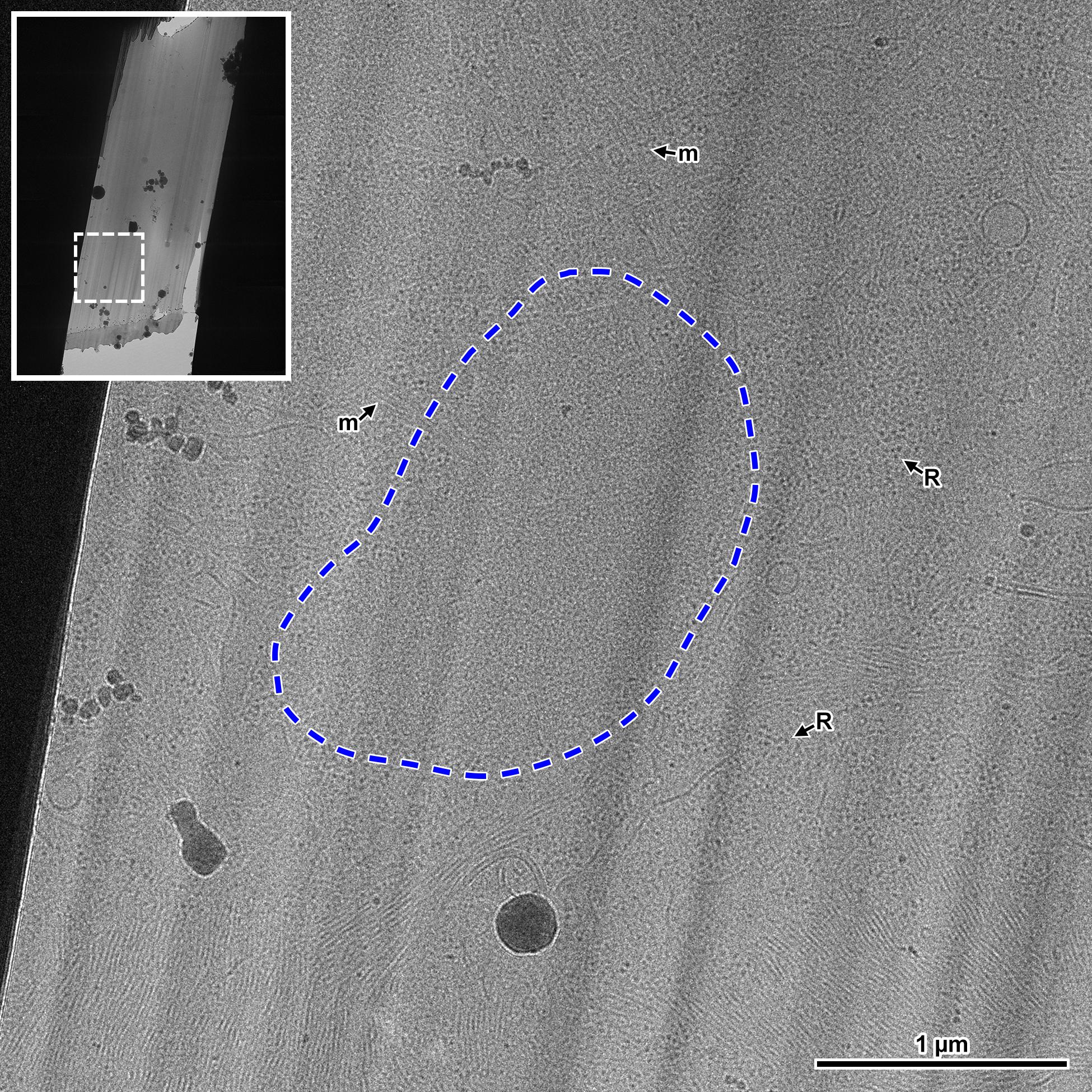

**Figure S28. Targeting images for chromatin in metaphase cells.**

Inset shows a montage of an entire lamella – the area bounded by the white dashed line corresponds to the location of the enlarged view. The region indicated by the blue dashed line contains chromatin and is shown at higher magnification as a cryotomographic slice in Figure 4C. Microtubules (m) and ribosomes (R) are also observed, and guided tilt-series target selection.

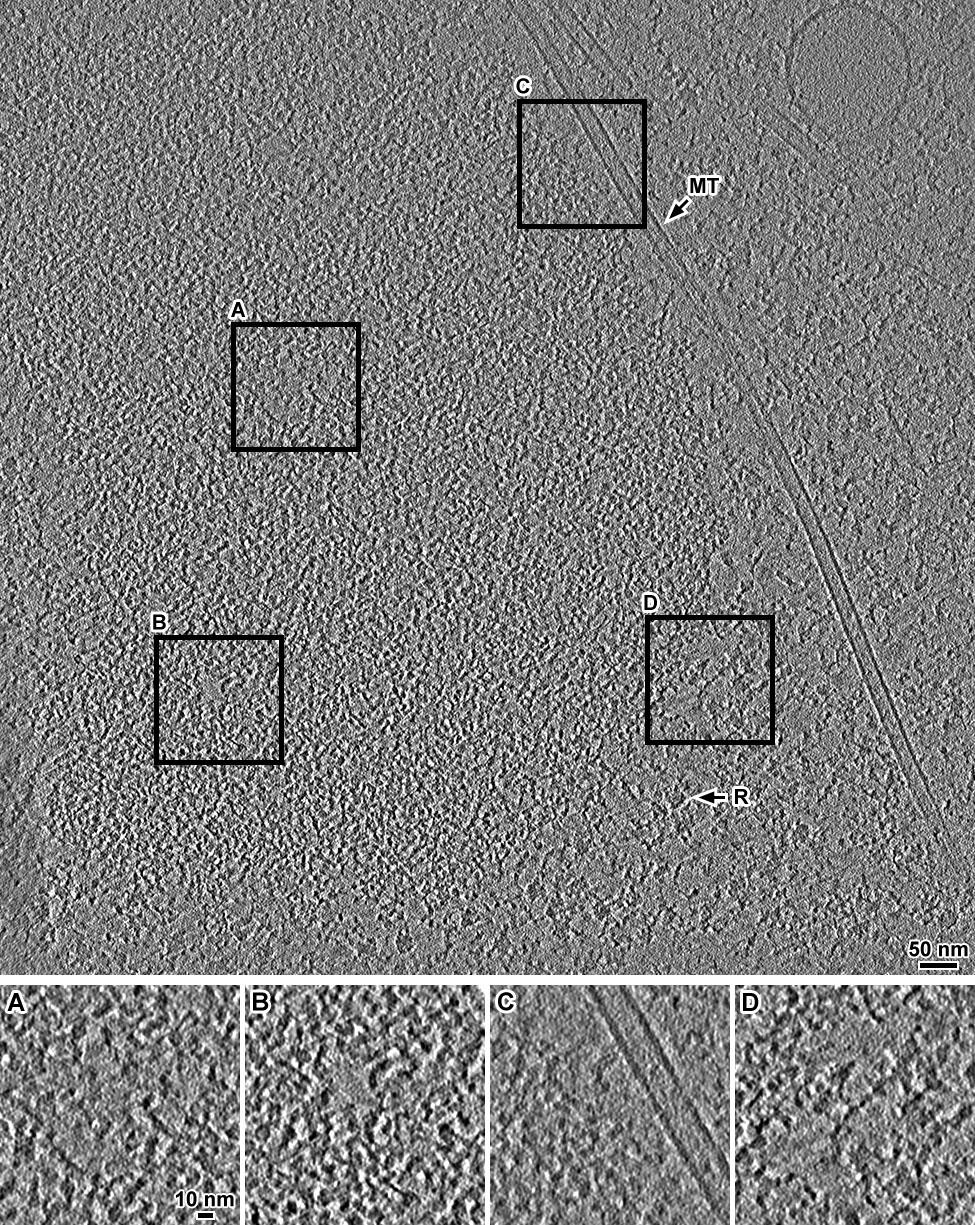

**Figure S29. Additional example Volta cryotomographic slice of a metaphase cell.**

Cryotomographic slice (20 nm) centered on a mitotic chromatin in a metaphase RPE-1 cell. Rendered with low JPEG compression. Insets show 2-fold enlargements of nucleosome-free pockets inside the chromosome (A and B) and the chromosome periphery (C and D). Cytological features are highlighted: ribosome (R); microtubule (MT).

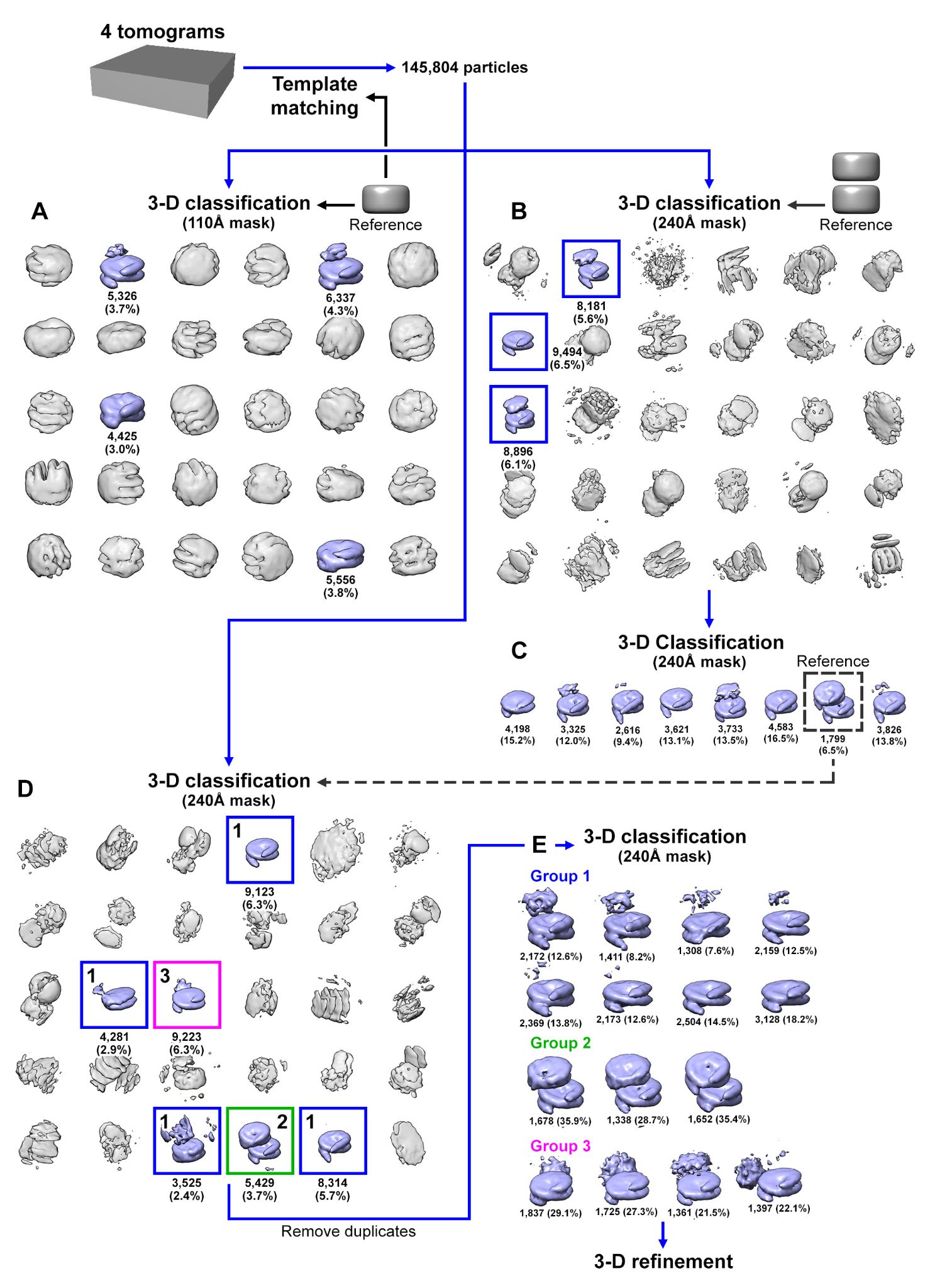

**Figure S30. Classification flowchart of M cells.**

(A) Metaphase candidate nucleosome subtomograms that were template matched with a cylindrical reference were directly classified in 3-D, using a cylindrical reference and a 120 Å spherical mask. (B) In parallel, the same set of subtomograms were directly classified in 3-D using a stacked cylinder reference and a larger spherical mask. The ambiguous densities (gray) are not canonical nucleosomes; they are abundant because the template matching process uses a featureless cylinder reference and a low cross-correlation cutoff. As a result, large numbers of false positives are rejected in the classification analysis. (C) A second round of classification using a nucleosome class average from B as the reference and the larger mask yielded mononucleosomes with extra densities at their face plus an unambiguous dinucleosome class. (D) Direct 3-D classification was done on the original set of subtomograms using a larger mask and the stacked dinucleosome class average from panel C as the reference. Three groups of class averages were obtained, corresponding to (1) mononucleosome, (2) stacked dinucleosome, and (3) mononucleosome with a gyre-proximal density. (E) These three groups of class averages were subjected to a third round of classification using either a mononucleosome, stacked dinucleosome or mononucleosome with a gyre-proximal density from panel D as the reference and large spherical mask.

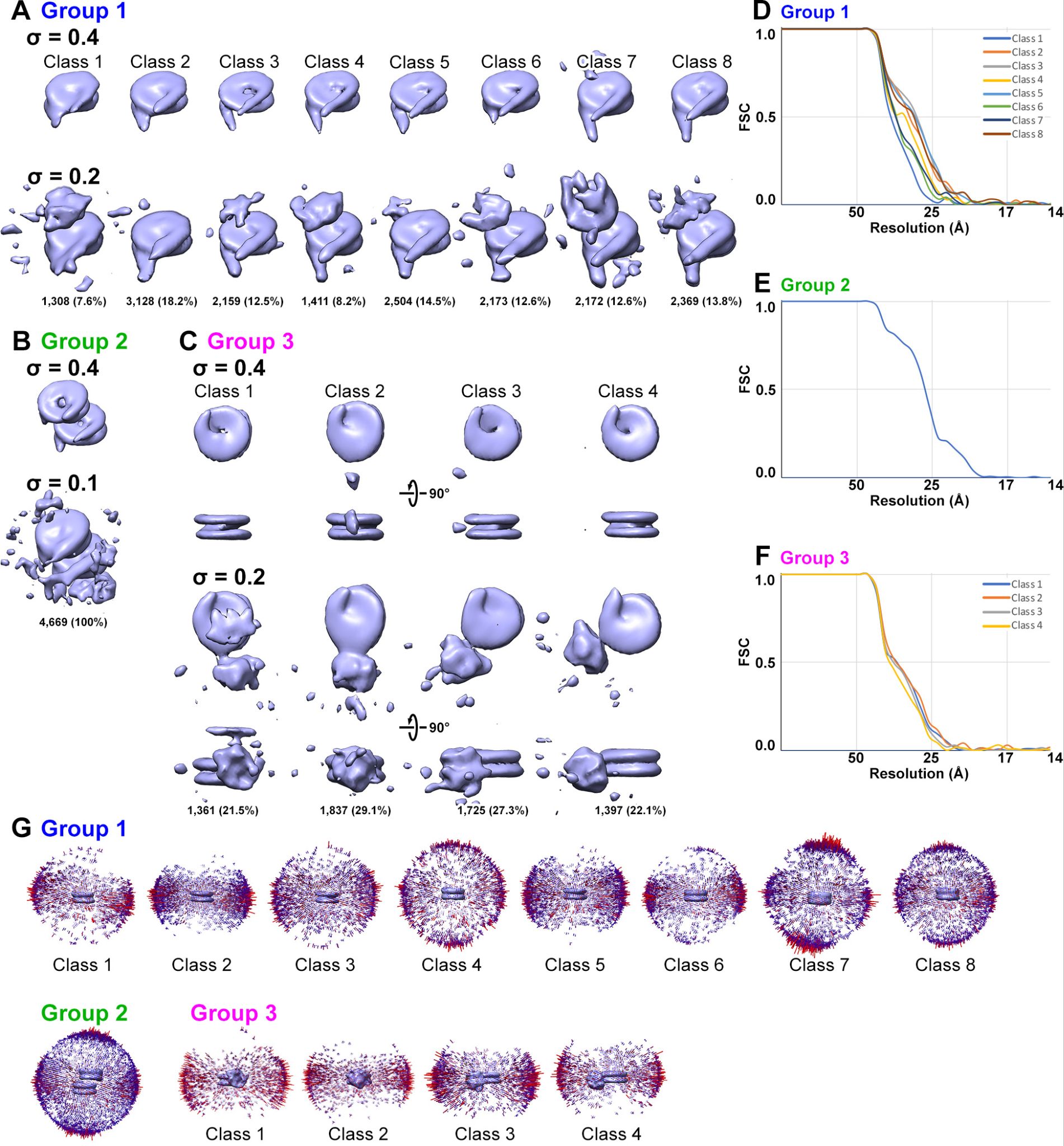

**Figure S31. Refinement of metaphase mononucleosomes and dinucleosomes.**

Refined class averages for (A) mononucleosomes, (B) stacked dinucleosomes, and (C) mononucleosomes with a gyre-proximal density. Two contour levels are shown for each class average. (D, E, F) FSC plots and (G) angular distribution of the refined density maps. For panels A – C, the hide dust feature in UCSF Chimera was disabled.

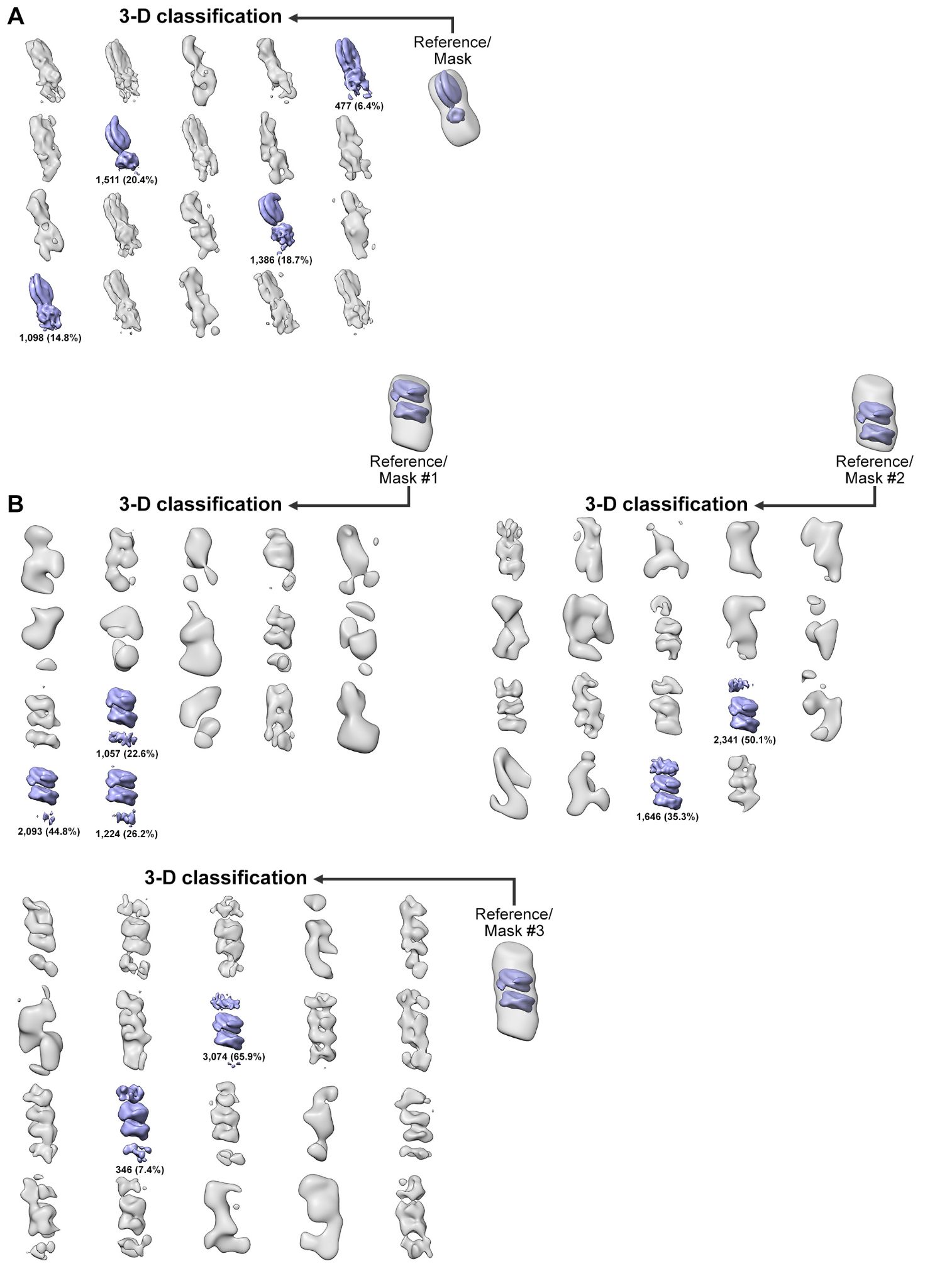

**Figure S32. Subtomogram analysis of alternative ordered nucleosome packing motifs in metaphase cells.**

Similar to Figure S16, nucleosome particles from Groups 2 and 3 (Figures S31 B and C, respectively) were subjected to an additional round of 3-D classification, using custom masks that enclose volumes where an additional complex may reside. Since the particles were already aligned from the previous refinement step, a restricted angular search range was imposed for these runs. The “reference/mask” models in the figure depicts the location of the volume masked-in (gray), relative to the reference (blue) used for each classification run. The masks used for these 3-D classification runs were optimized for (A) side-by-side nucleosomes and (B) ordered trinucleosomes. Class averages that contain at least one ordered nucleosome are shaded blue.

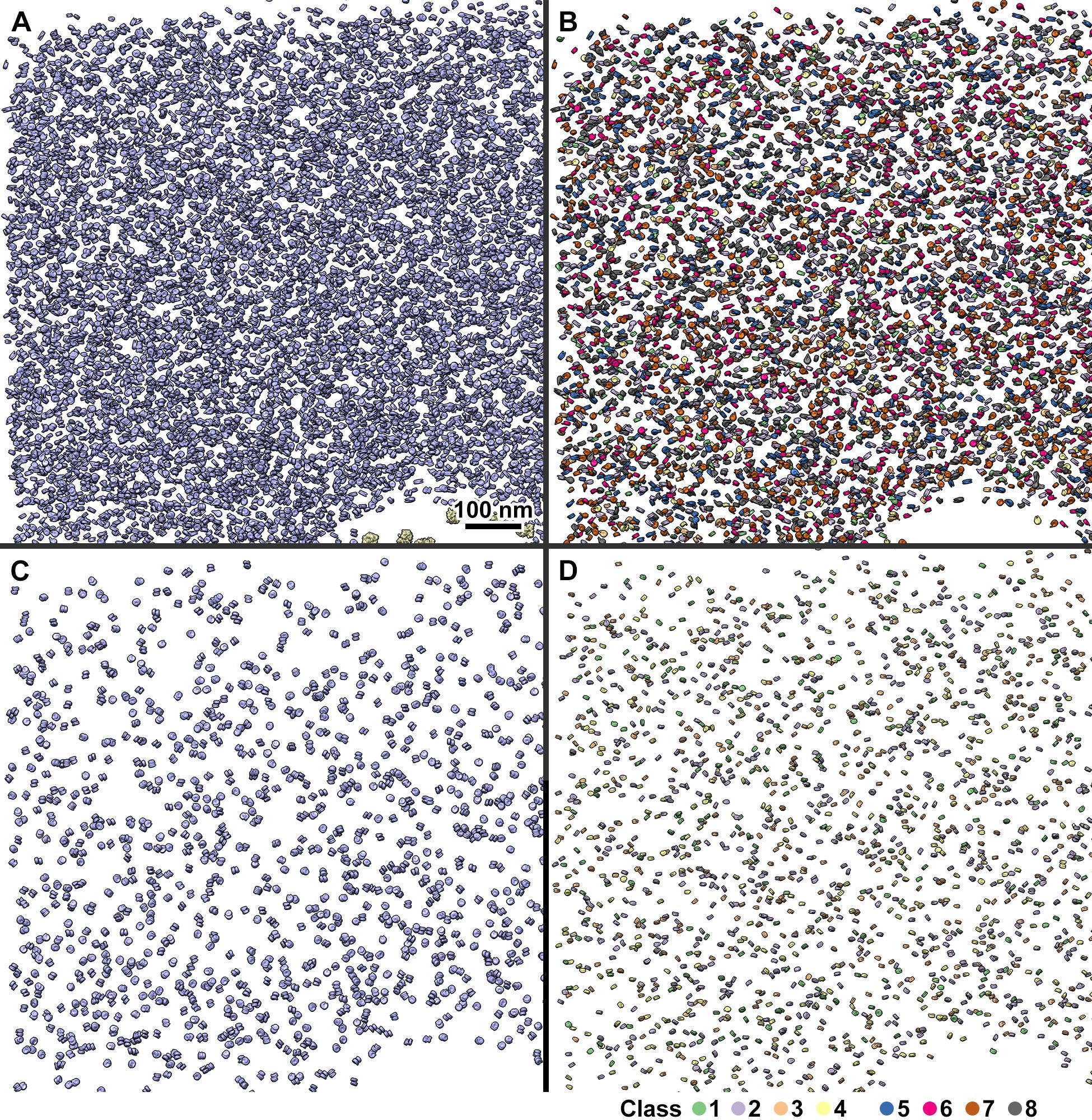

**Figure S33. Remapped models of metaphase nucleosome groups.**

(A) Enlargement of a segment of Figure 5D showing all remapped nucleosomes (blue) in a G1 domain. Class averages of mononucleosomes (group 1), ordered stacked dinucleosomes (group 2) and mononucleosomes with gyre-proximal density (group 3) are remapped separately in panels B, C and D respectively. The color-coding of the nucleosome densities in panels B and D correspond to the class averages shown in Figure 5A and Figure 5C.

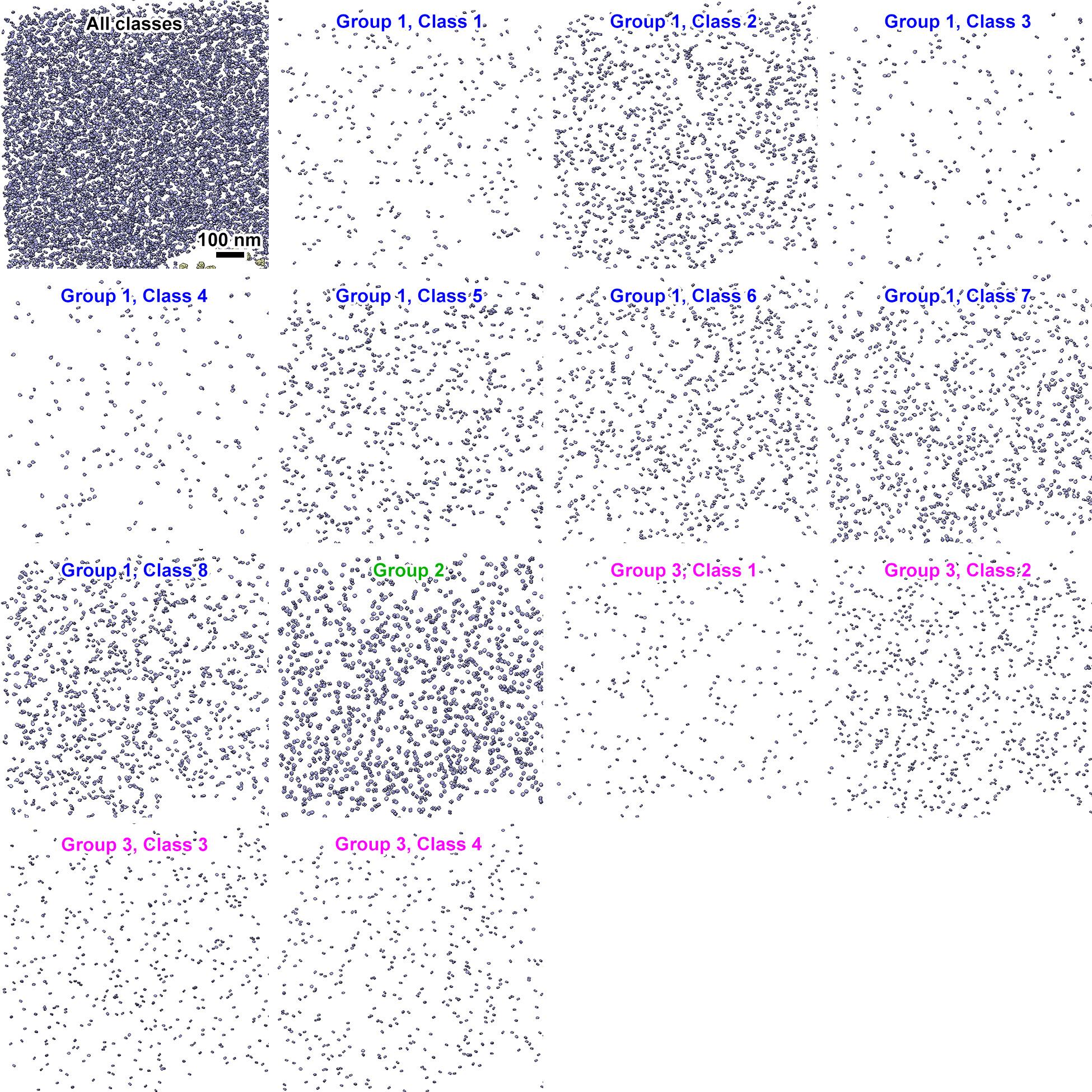

**Figure S34. Remapped models of metaphase nucleosome individual classes.**

The upper left panel shows a section of the remapped model presented in Figure 5D. The remaining panels also show the same section of the remapped model, but with each of the nucleosome class averages from Figure 5A – C rendered separately.

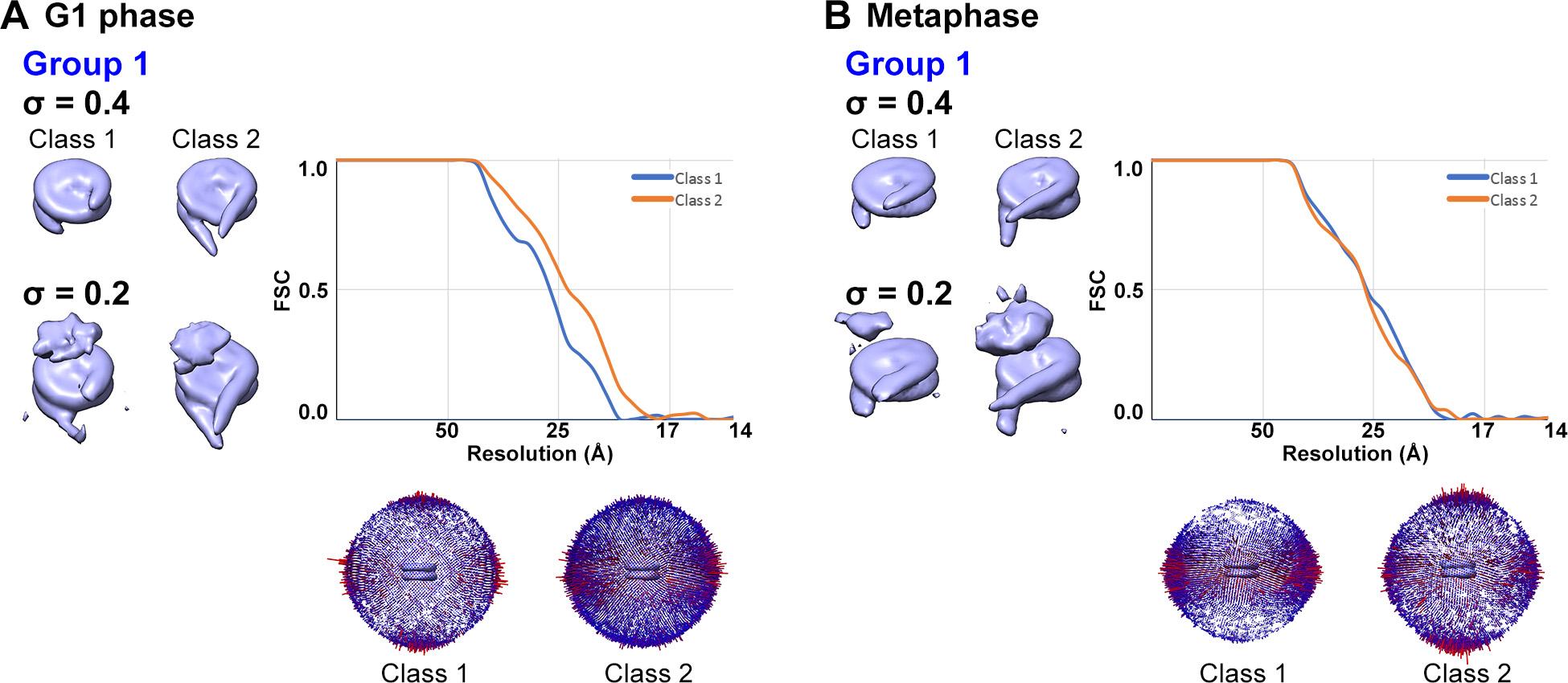

**Figure S35. Refinement of Group 1 mononucleosomes combined into two classes.**

Mononucleosome classes from Group 1 (Figures 3A and 5A) were manually assigned into two groups, based on the length of the linker DNA. Shown here are the refinement results for the short linker (Class 1) and long linker (Class 2) class for (A) G1 and (B) metaphase cells, respectively. The refined mononucleosome class averages here were used for docking, as shown in Figure 7.

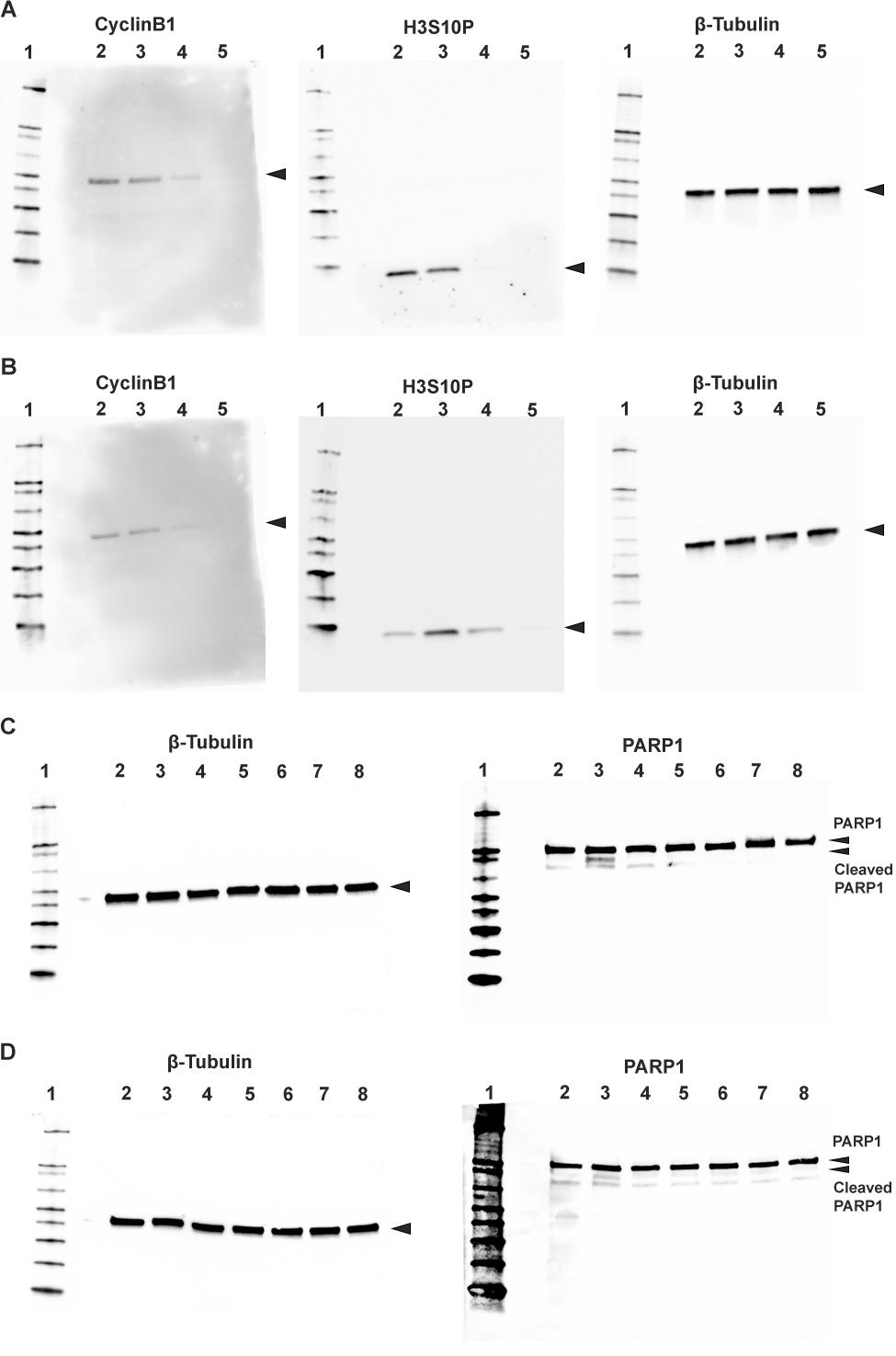

**Figure S36. Uncropped immunoblots.**

(A and B) Uncropped immunoblots from Figure S1. (A) Nocodazole arrest and washout. (B) MG132 arrest and washout. The lane number is indicated above the gels and correspond, respectively, to the protein ladder (Invitrogen, #LC5602); arrested cells; 30 minutes after washout; 60 minutes after washout; 120 minutes after washout.

(C) Apoptosis detection in G1 phase RPE1 cells treated with or without cryoprotectant. (D) Apoptosis detection in metaphase RPE1 cells treated with or without cryoprotectants. Lanes 1, 2, 3 in both (C) and (D) correspond respectively to the protein ladder, 0.1% DMSO treatment for 6 hours, and 1 μM staurosporine treatment for 6 hours. Lanes 4, 5, 6, 7, 8 in (C) are G1 phase RPE1 cells, the lanes correspond respectively to the G1 cells, G1 cells treated with 9% DMSO for 1 minute, G1 cells treated with 9% DMSO for 10 minutes, G1 cells treated with 9% glycerol for 1 minute, G1 cells treated with 9% glycerol for 10 minutes. Lanes 4, 5, 6, 7, 8 in (D) are metaphase RPE1 cells, the lanes correspond respectively to the metaphase cells, metaphase cells treated with 9% DMSO for 1 minute, metaphase cells treated with 9% DMSO for 10 minutes, metaphase cells treated with 9% glycerol for 1 minute, metaphase cells treated with 9% glycerol for 10 minutes. The positions of PARP1 and cleaved PARP1 proteins are indicated with the arrowheads.

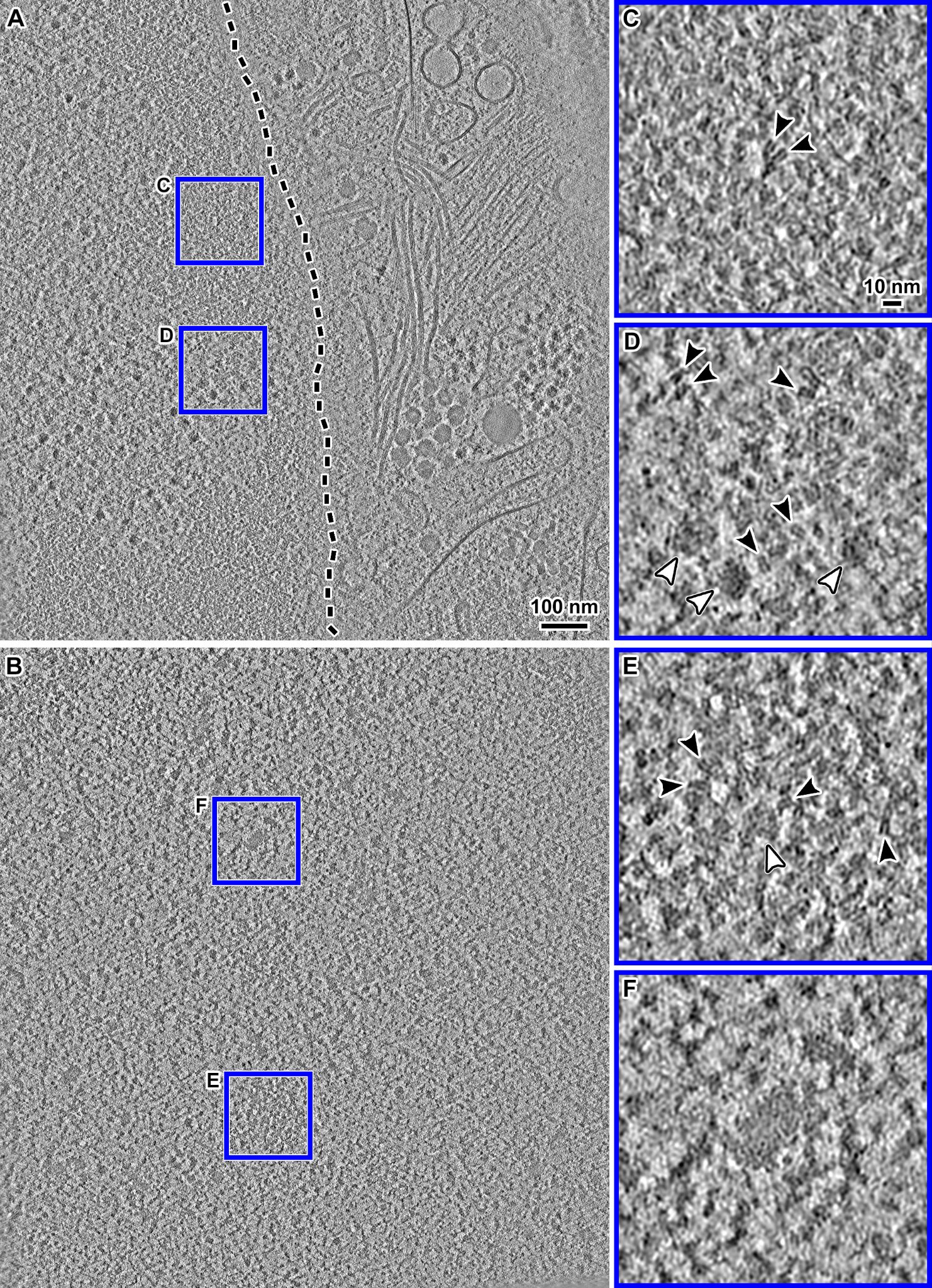

**Figure S37. Non-denoised version of Figure 2.**

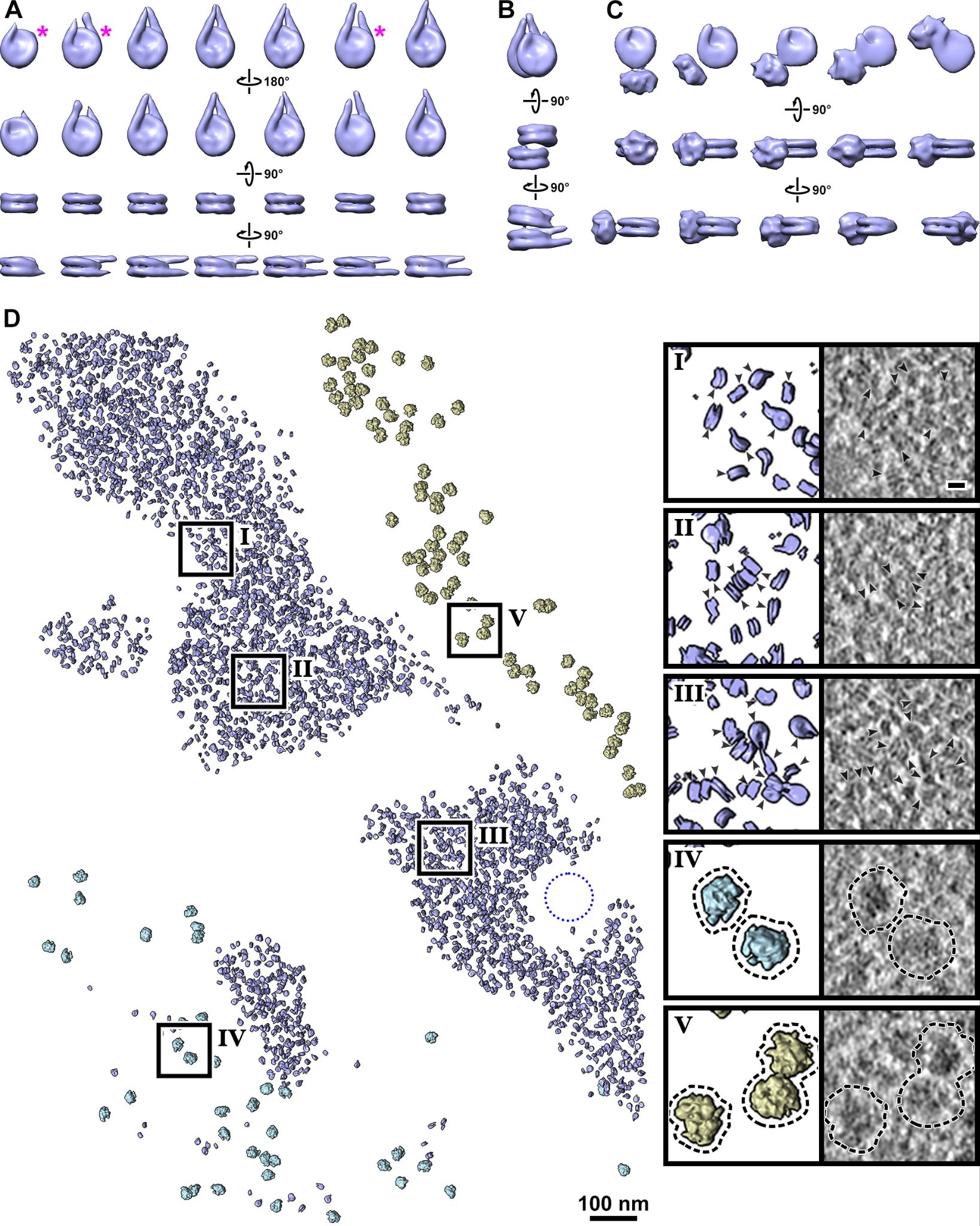

**Figure S38. Reproduction of Figure 3, but with non-denoised versions of the cryotomographic slices.**

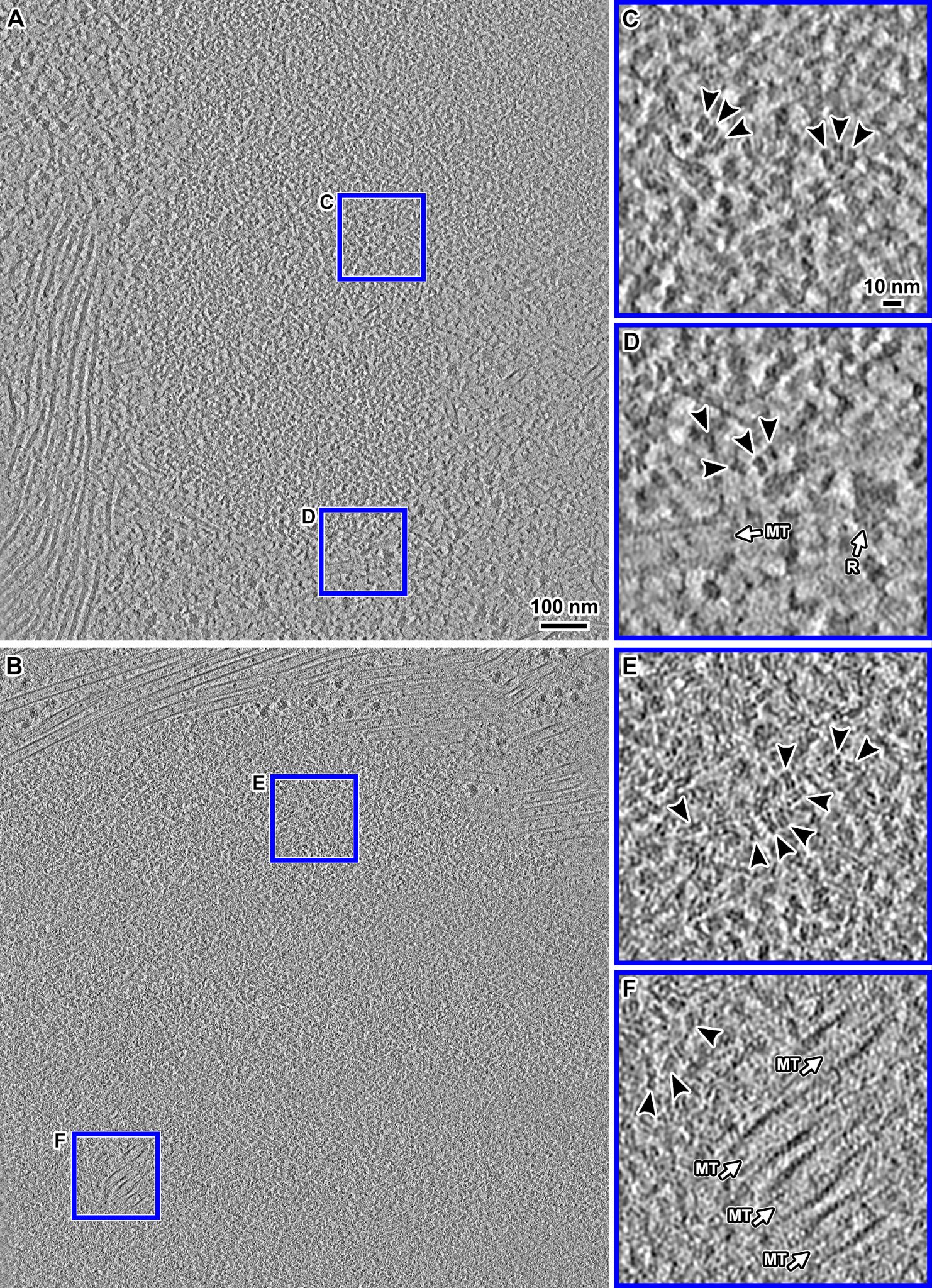

**Figure S39. Non-denoised version of Figure 4.**

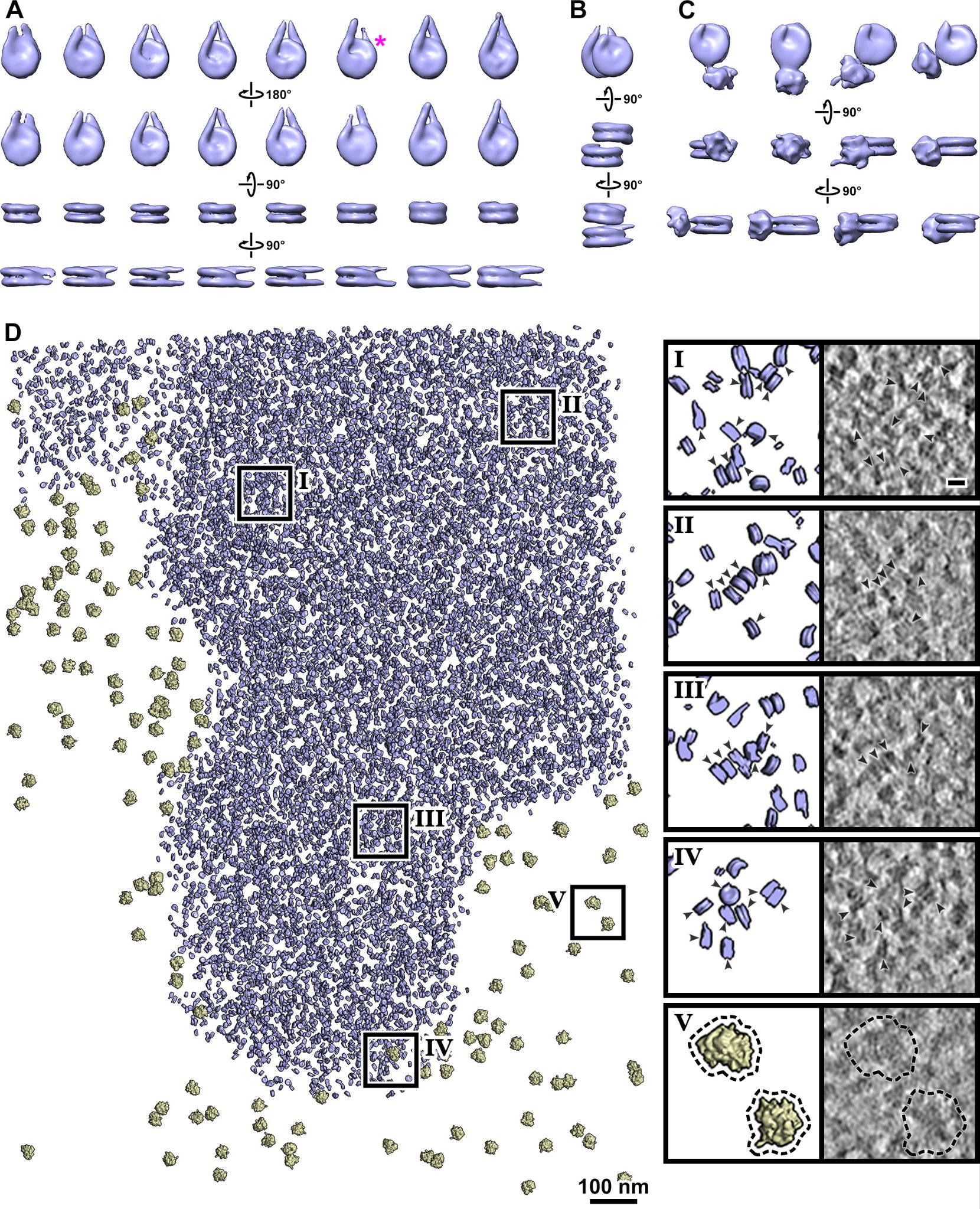

**Figure S40. Reproduction of Figure 5, but with non-denoised versions of the cryotomographic slices.**
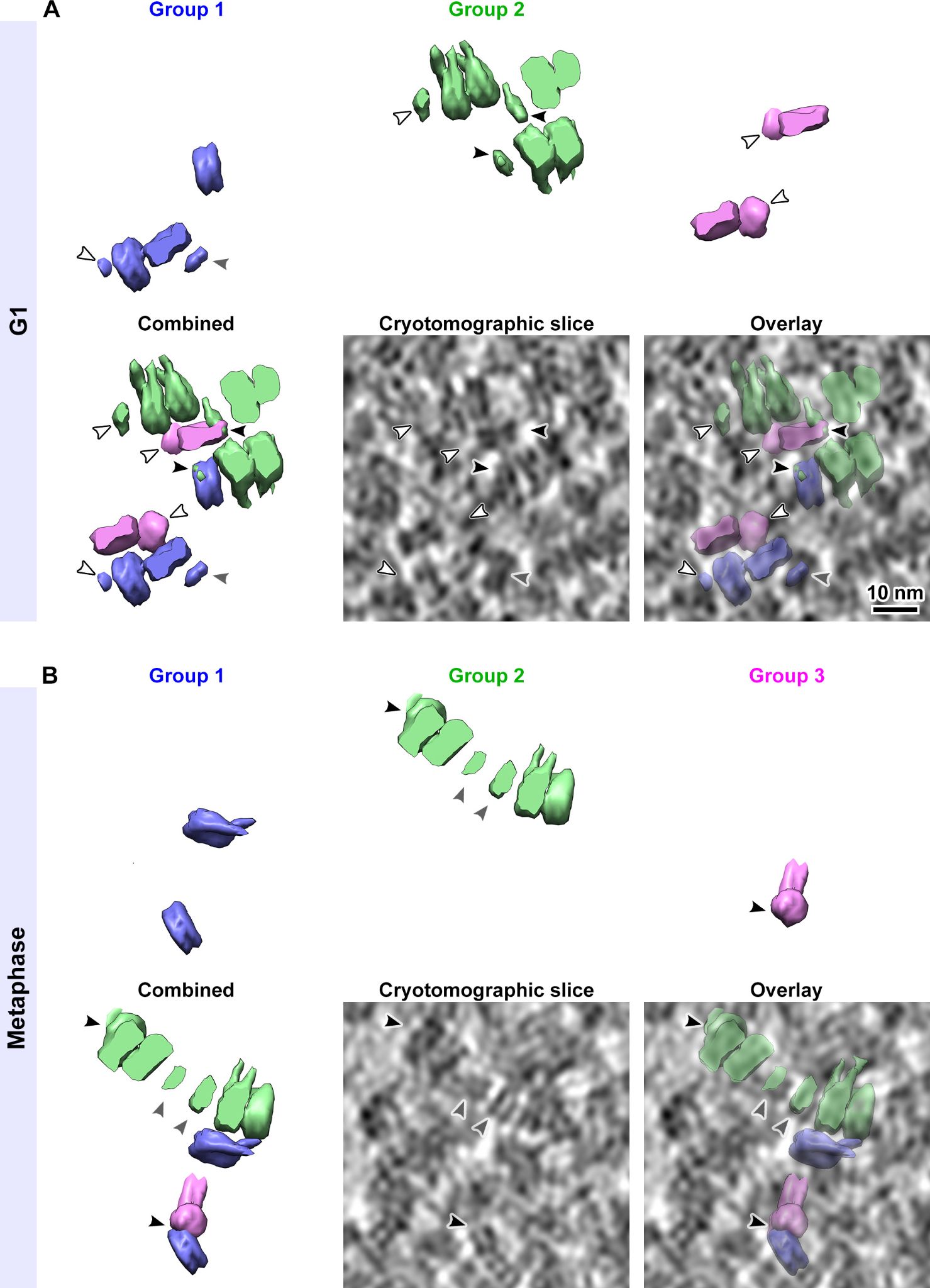

**Figure S41. Reproduction of Figure 6, but with non-denoised versions of the cryotomographic slices.**

**Movie S1. Subtomogram averages of nucleosomes in G1 cell nuclei.**

Subtomogram averages of mononucleosomes, stacked dinucleosomes, and nucleosomes with a gyre-proximal density, all from G1 chromatin domains. See also Figure 3, A – C.

**Movie S2. *In situ* overview of G1 cell nucleus.**

The cryotomogram is rendered as 10 nm slices. Regions highlighted in blue indicate chromatin domains that were segmented using EMAN2. The remapped model shows nucleosomes (blue) and preribosomes (light blue) in the nucleus and ribosomes (yellow) in the cytoplasm. See also Figure 3D, which is rotated 90° counterclockwise relative to this movie.

**Movie S3. Subtomogram averages of nucleosomes in metaphase chromatin.**

Subtomogram averages of mononucleosomes, stacked dinucleosomes, and nucleosomes with a gyre-proximal density, all in metaphase chromatin. See also Figure 5, A – C.

**Movie S4. *In situ* overview of metaphase chromatin.**

The cryotomogram is rendered as 10 nm slices. The region highlighted in blue indicates chromatin, segmented using EMAN2. The remapped model shows nucleosomes (blue) and ribosomes (yellow). See also Figure 5D, which is rotated 90° counterclockwise relative to this movie.

**Table 1. Relative abundance of ordered nucleosome species.**

|  | **NCP** | **Di-NCP** | **Proximal** |
| --- | --- | --- | --- |
| **G1 phase** | 1.00 | 0.22 | 0.49 |
| **Metaphase** | 1.00 | 0.27 | 0.37 |

NCP = mononucleosomes (all linker DNA lengths), Di-NCP = stacked dinucleosomes, Proximal = nucleosomes with proximal density. The ratio is normalized to the number of NCP particles within each cell-cycle state.

**Table 2. Summary of chromatin and nuclear features *in situ*.**

| **Structural feature** | **RPE-1 G1** | **RPE-1 Meta** | **HeLa †** | **Yeast †** |
| --- | --- | --- | --- | --- |
| **≥ 100 nm** |  |  |  |  |
| Mitotic chromatin plates | n/a | Not seen | n/a | Not seen |
| Arc-shaped stacks / slinkies | Not seen | Not seen | Not seen | Not seen |
| 100 – 200 nm periodic structure | Not seen | Not seen | Not seen | Not seen |
| Interphase domain shape | Irregular | n/a | Irregular | n/a |
| **20 – 80 nm** |  |  |  |  |
| Chromatin packing | Irregular | Irregular | Irregular | Irregular |
| Ordered trinucleosome stacks | Not seen | Not seen | Not seen | Not seen |
| Slinky oligonucleosomes | Not seen | Not seen | Not seen | Not seen |
| Megacomplexes (in chromatin) | Rare | Rare | Rare | n/a |
| Megacomplexes (in nucleoplasm) | Abundant | n/a | Abundant | Abundant |
| Dense irregular bodies | Rare | Not seen | Rare | in *S. cere* |
| Pockets (mitotic chromatin) | n/a | Rare | n/a | in *S. pombe* |
| Pockets (G1 chromatin domain) | Ambiguous | n/a | n/a | n/a |
| **10 – 20 nm** |  |  |  |  |
| Ordered dinucleosome stacks * | Abundant | Abundant | Not seen | Not seen |
| Gyre-interacting densities * | Abundant | Abundant | n/d | n/d |
| Canonical mononucleosome | Abundant | Abundant | Abundant | Rare |
| **< 10 nm** |  |  |  |  |
| Ordered linker DNA length | Variable | Variable / no ultrashort | Long & short | Long & short |
| Linker DNA length symmetry | Asymmetric & symmetric | Asymmetric & symmetric | Asymmetric | Asymmetric |

The chromatin and nuclear structural features (approximate size ranges in nanometers) of RPE-1 G1 phase (G1), Metaphase (Meta), HeLa interphase, and *S. pombe* and *S. cerevisiae* in interphase and mitosis (Yeast) are listed. † The HeLa phenotypes are from (Cai *et al*, 2018a) while the yeast phenotypes are from (Cai *et al*, 2018b; Chen *et al*, 2016; Ng *et al*, 2019; Tan *et al*, 2023); note that these earlier studies have a smaller sample size of canonical nucleosomes than here. “Not seen” means that the expected structures were absent in our dataset. We cannot rule out that they exist at low abundance. n/a: not applicable. n/d: not determined due to sample size issues. * Some ordered higher-order nucleosome structures may be sensitive to cryoprotectants or image contrast effects.

**Table S1. Research resources.**

| **Resource** | **Source** | **Catalog ID / link** |
| --- | --- | --- |
| **Chemicals** |  |  |
| 4′,6-diamidino-2-phenylindole (DAPI) | TFS | D1306 |
| Dimethyl sulfoxide (DMSO) | Sigma | 472301 |
| DMEM/F-12 with GlutaMAX | Gibco/TFS | 10565-018 |
| Fetal bovine serum | Sigma | F9665 |
| Glycine | Bio-Rad | 1610718 |
| MG132 | Merck | M7449-1ML |
| Nocodazole | Merck | SML1665-1ML |
| Palbociclib | Selleckchem | S1116 |
| Paraformaldehyde | EMS | 15714 |
| Penicillin-streptomycin | Gibco | 15140-122 |
| Phosphate buffered saline (PBS) | Vivantis | PB0344-1L |
| PBS-T (PBS + 0.1% Tween 20) | Sinopharm | T20087687 |
| Prolong Gold antifade reagent | TFS | P36930 |
| Taxol | Santa Cruz | 33069-62-4 |
| Triton X-100 | Alfa Aesar | A16046 |
| **Antibodies** |  |  |
| Alexa Fluor® 647 anti-histone H3 (phospho S10) | Abcam | ab196698 |
| Rabbit polyclonal Anti-phospho-Histone H3 (Ser10) | Abcam | ab5176 |
| Mouse anti-rabbit IgG-HRP | Santa Cruz | sc-2357 |
| Rabbit polyclonal to beta Tubulin | Abcam | ab6046 |
| m-IgGκ BP-HRP | Santa Cruz | sc-516102 |
| cyclin B1 (GNS1) mouse monoclonal IgG1 | Santa Cruz | sc-245 |
| Anti-RPB1 CTD(S5P) | Abcam | ab5131 |
| Anti-RPB1 CTD(S2P) | Abcam | ab5095 |
| **Cell line** |  |  |
| hTERT-RPE-1 cells | ATCC | CRL-4000 |
| **Chromatin** |  |  |
| HeLa oligonucleosomes | EpiCypher | 16-0003 |
| **Light microscopy** |  |  |
| LSM900 AiryScan microscope | Zeiss | N/A |
| 20×, 0.8 N.A. Plan-Apochromat | Zeiss | N/A |
| 63×, 1.4 N.A. oil Plan-Apochromat | Zeiss | N/A |
| ZEN 3.0 (blue edition) | Zeiss | N/A |
| Eclipse Ti microscope | Nikon | N/A |
| Plan Fluor 10x Ph1 objective | Nikon | N/A |
| 20 mm Ø coverslip | Paul Marienfeld GmbH | 0112600 |
| **Electron cryomicroscopy** |  |  |
| C-flat 2/4 200 mesh, gold | Protochips | CF-2/4-2Au |
| C-Clip Ring (autogrid) | TFS | 1036173 |
| CryoFIB autogrid | TFS | 1205101 |
| C-flat 2/4 200 mesh, copper | Protochips | CF-2/4-2C |
| Continuous carbon | EMS | CF200-Cu-UL |
| Whatman Grade 1 filter paper | Whatman | 1001-055 |
| Copper tubes, 0.3mm inner diameter | Wohlwend | N/A |
| Copper tube loading tool | Wohlwend | Part 733-1 |
| Copper tube cutting device | Wohlwend | Part 732 |
| UC7/FC7 Microtome | Leica | N/A |
| 35° diamond knife | Diatome | Cryo35 |
| Micromanipulator, Leica | Leica | N/A |
| Micromanipulator, MN-151-S | Narishige | N/A |
| Vitrobot Mark IV | TFS | N/A |
| Helios Nanolab 650 FIB-SEM | TFS | N/A |
| PolarPrep 2000 Cryo Transfer System | Quorum | N/A |
| Titan Krios G1 cryo-TEM | TFS | N/A |
| Falcon II camera | TFS | N/A |
| Gatan K3 Summit camera | AMETEK | N/A |
| Gatan BioContinuum imaging filter | AMETEK | N/A |
| **Software** |  |  |
| Adobe Illustrator | Adobe | adobe.com |
| Adobe Photoshop | Adobe | adobe.com |
| Adobe Premiere Pro | Adobe | adobe.com |
| Auxiliary scripts | Gan lab | github.com/anaphaze/ot-tools |
| BioRender | BioRender | biorender.com |
| Bsoft | (Heymann & Belnap, 2007) | lsbr.niams.nih.gov/bsoft |
| EMAN2 | (Chen *et al*, 2017) | blake.bcm.edu/emanwiki/EMAN2 |
| FIJI | (Schindelin *et al*, 2012) | imagej.net/software/fiji |
| Google sheets | Google | www.google.com/sheets |
| IMOD | (Mastronarde, 1997) | bio3d.colorado.edu/imod |
| PEET | (Nicastro *et al*, 2006) | bio3d.colorado.edu/PEET |
| RELION | (Scheres, 2012) | github.com/3dem/relion |
| SerialEM | (Mastronarde, 2005) | bio3d.colorado.edu/SerialEM |
| UCSF Chimera | (Pettersen *et al*, 2004) | www.cgl.ucsf.edu/chimera |

**Table S2. Antibodies used.**

| **Antigen** | **1º antibody or**  **conjugated antibody** | **2º antibody** | **Dilution** | |
| --- | --- | --- | --- | --- |
|  |  |  | **1º** | **2º** |
| **Immunoblots** | | | | |
| H3S10P | Rabbit polyclonal Anti-phospho-Histone H3 (Ser10) Antibody (Abcam ab5176) | mouse anti-rabbit IgG-HRP (Santa Cruz sc-2357) | 1:500 | 1:5000 |
| Cyclin B | cyclin B1 (GNS1) mouse monoclonal IgG1 (Santa Cruz sc-245) | m-IgGκ BP-HRP  (Santa Cruz sc-516102) | 1:500 | 1:5000 |
| Beta Tubulin | Rabbit polyclonal to beta Tubulin (Abcam ab6046) | mouse anti-rabbit IgG-HRP (Santa Cruz sc-2357) | 1:500 | 1:5000 |
| PARP1 | Rabbit monoclonal [EPR18461] to PARP1 | mouse anti-rabbit IgG-HRP (Santa Cruz sc-2357) | 1:500 | 1:5000 |
| **Immunofluorescence** | | | | |
| H3S10P | Alexa Fluor 647 Mouse Monoclonal Anti-Histone H3 (phospho S10) (Abcam ab196698) | N/A | 1:250 | N/A |
| RPB1 CTD(S5P) | Rabbit polyclonal, (Abcam ab5131) | Goat anti-Rabbit IgG (H+L) Secondary Antibody, Alexa Fluor 488 (TFS A-11008) | 1:300 | 1:500 |
| RPB1 CTD(S2P) | Rabbit polyclonal, (Abcam ab5095) | Goat anti-Rabbit IgG (H+L) Secondary Antibody, Alexa Fluor 488 (TFS A-11008) | 1:300 | 1:500 |

TFS = Thermo Fisher Scientific.

**Table S3. Confocal microscopy details.**

| **General** |  |
| --- | --- |
| Model | Zeiss LSM900 |
| Control software | cellSense v X.Y |
| Pinhole | 1 AU |
| X, Y pixel | 0.087 [µm] |
| Z pixel | 0.410 [µm] |
| **Acquisition** |  |
| Objective Lens | UPLSAPO 60XO |
| Objective Lens Mag. | 60× |
| Objective Lens NA | 1.35 |
| Scan Device | Galvano |
| Scan Direction | One way |
| Sampling Speed | 2.0 [µs/pixel] |
| Sequential Mode | Line |
| Integration Type | None |
| Integration Count | 0 |
| Zoom | ×2.38 |
| **GFP channel settings** |  |
| Emission WaveLength | 510 [nm] |
| PMT Voltage | 500 [V] |
| C.A. | 200 [µm] |
| Bits/Pixel | 12 [bits] |
| Laser Wavelength | 488 [nm] |
| Laser Transmissivity | 0.05 [%] |
| AOTF/AOM Transmissivity | 0.5 [%] |
| Laser ND Filter | 10 [%] |
| Detection Wavelength | 500 – 600 [nm] |
| **DIC channel settings** |  |
| PMT Voltage | 380 [V] |
| C.A. | 200 [µm] |
| Bits/Pixel | 12 [bits] |
| Laser Wavelength | 488 [nm] |
| Laser Transmissivity | 0.05 [%] |
| AOTF/AOM Transmissivity | 0.5 [%] |
| Laser ND Filter | 10 [%] |
| **Live cell imaging** |  |
| Interval | 30 seconds |
| Duration | 2 hours |
| FoV | 50 µm × 50 µm |

**Table S4. Cryo-ET imaging details.**

| **Sample preparation** |  |
| --- | --- |
| EM grids | Cryosections: continuous carbon  Lamellae: gold |
| Plunge freezer | Vitrobot Mark IV |
| Cryomicrotome | UC7/FC7 |
| Attachment device | Crion |
| Micromanipulators | Leica micromanipulator; Narishige MN-151-S |
| Cryomicrotome feed | 100 nm |
| Cryo-FIB-SEM | Helios NanoLab 650 DualBeam |
| Cryo-transfer device | Quorum PP2000T |
| Milling currents | Rough: 2.8 nA  Intermediate: 0.28 nA  Polishing: 48 pA |
| **Cryo-ET data collection** |  |
| Microscope | Titan Krios |
| Energy | 300 keV |
| Camera: recording mode | FII: integration  K3: super-resolution movie frames |
| Energy filter width | K3: 20 eV |
| Tomography software | TFS Tomo4 and SerialEM |
| Unbinned pixel size | FII: 7.3, 12.4Å  K3: 3.4, 4.5 Å |
| Contrast mechanism | Defocus phase contrast (cryolamellae, cryosections, oligonucleosomes)  Volta phase contrast (cryolamellae, cryosections) |
| Defocus (nominal) | Defocus phase contrast: −5 (K3) or −10 µm (FII)  Volta phase contrast: −0.5 μm |
| Cumulative dose | cryosections: 100 e^−^ / Å^2^  oligonucleosomes: 120 e^−^ / Å^2^  cryolamellae: 110 e^−^ / Å^2^ |
| Dose fractionation | cryosections: 1/cosine  oligonucleosomes and cryolamellae: (1/cosine)^(1/y), where y = 4 |
| Tilt range | cryosections: ±60°; bidirectional, negative angles first  cryolamellae: −70° to +42° start −14°, dose-symmetric  oligonucleosomes: ±60° start 0°; dose-symmetric, negative angles first |
| Tilt increment | 2° |
| **Cryo-ET data analysis** |  |
| Tomogram processing | IMOD 4.11 |
| Template matching | PEET 1.15 |
| Reference creation | Bsoft 1.8.8 |
| Mask creation | Bsoft 1.8.8, RELION 3.0.8 |
| Subtomogram analysis | RELION 3.0.8, Auxiliary scripts |
| Tomogram visualization | UCSF Chimera 1.15, IMOD 4.11 |
| Auxiliary scripts | github.com/anaphaze/ot-tools |
| Calculations | Google sheets, FIJI |
| Figure/movie editing | Adobe Photoshop, Illustrator, and Premiere Pro, www.biobender.com |

**Table S5. Cryotomogram details.**

| **Tomogram** | **Description** | **Sample** | **Fig** | **Dose (e/Å^2^)** | **Pixel size (Å)** | **∆f ***  **(μm)** | **Cam** | **VPP** | **thick (nm)** | **resid** |
| --- | --- | --- | --- | --- | --- | --- | --- | --- | --- | --- |
| 20221029_03 | G1, 9% DMSO | lamella | 2 | 110 | 3.4 | 0.5 | K3 | + | 70 | 0.49 |
| 20221028_39 | G1, 9% DMSO | lamella | 2 | 110 | 3.4 | 0.5 | K3 | + | 190 | 0.57 |
| 20221028_14 | G1, 9% DMSO | lamella | 3, 6, S23, Mov S2 | 110 | 3.4 | 0.5 | K3 | + | 80 | 0.57 |
| 20221028_[10, 18, 19, 21, 23, 24, 29, 30, 41]  20221029_[20, 32, 40, 49, 51, 53]  20221030_[36, 41] | G1, 9% DMSO | lamella | - | 110 | 3.4 | 0.5 | K3 | + | 70– 210 | 0.31– 1.40 |
| 20220809_05 | M, 9% DMSO | lamella | 4, 6 | 110 | 3.4 | 0.5 | K3 | + | 150 | 0.31 |
| 20230209_28 | M, 9% DMSO | lamella | 4, Mov S4 | 110 | 3.4 | 0.5 | K3 | + | 110 | 0.56 |
| 20230209_[42, 58] | M, 9% DMSO | lamella | - | 110 | 3.4 | 0.5 | K3 | + | 120, 80 | 0.85, 0.90 |
| 20161115_008 | M | section | S2 | 95 | 12.4 | 11.1 | FII | − | 160 | 0.56 |
| 20180528_02 | M | section | S2 | 100 | 7.3 | 0.5 | FII | + | 170 | 0.86 |
| 20201217_01 | G1, 0% DMSO | lamella | S4A | 120 | 3.4 | 7.0 | K3 | − | 140 | 0.76 |
| 20210110_20 | G1, 3% DMSO | lamella | S4B | 110 | 3.4 | 8.1 | K3 | − | 120 | 0.20 |
| 20210303_16 | G1, 6% DMSO | lamella | S4C | 110 | 3.4 | 8.4 | K3 | − | 200 | 0.52 |
| 20210303_02 | G1, 9% DMSO | lamella | S4D | 110 | 3.4 | 8.2 | K3 | − | 110 | 0.79 |
| 20210423_17 | M, 9% DMSO | lamella | S5A | 110 | 3.4 | 4.9 | K3 | − | 150 | 1.29 |
| 20210526_01 | M, 9% Glycerol | lamella | S5B | 110 | 3.4 | 4.2 | K3 | − | 160 | 0.54 |
| 20210923_03 | M, 9% DMSO | lamella | S5C | 110 | 3.4 | 0.5 | K3 | + | 110 | 0.91 |
| 20211101_16 | M, 9% Glycerol | lamella | S5D | 110 | 3.4 | 0.5 | K3 | + | 180 | 0.73 |
| 20210916_03 | Oligo | plunge | S6A | 120 | 3.4 | 0.5 | K3 | + | ~90 | 0.27 |
| 20210904_26 | Oligo, 9% DMSO | plunge | S6B | 120 | 3.4 | 0.5 | K3 | + | ~90 | 0.41 |
| 20221028_31 | G1, 9% DMSO | lamella | S10 | 110 | 3.4 | 0.5 | K3 | + | 210 | 0.31 |
| 20210923_09 | M, 9% DMSO | lamella | S29 | 110 | 3.4 | 0.5 | K3 | + | 180 | 0.41 |
| 20210916_[05, 06, 07, 09–33] | Oligo | plunge | - | 120 | 3.4 | 0.5 | K3 | + | ~90 | 0.27–1.04 |
| 20210904_[12–41}] | Oligo, 9% DMSO | plunge | - | 120 | 3.4 | 0.5 | K3 | + | ~90 | 0.35–1.39 |
| 20240101_[29, 53] | G1, 9% Glycerol | lamella | S17 | 110 | 3.4 | 0.5 | K3 | + | ~90, 70 | 0.26, 0.29 |
| 20240101_[21, 31, 44] | G1, 9% Glycerol | lamella | - | 110 | 3.4 | 0.5 | K3 | + | 90– 120 | 0.29– 0.62 |
| 20231207_[25, 27] | G1 | lamella | S20 | 110 | 3.4 | 0.5 | K3 | + | 80, 110 | 0.27, 0.30 |
| 20231207_[33, 37, 38, 40, 42], 20231215_[24, 26] | G1 | lamella | - | 110 | 3.4 | 0.5 | K3 | + | 70– 160 | 0.28– 0.52 |

The tilt increment was 2° for all sets. All data reported in this table were used for subtomogram analysis and were deposited as EMPIAR-#####. Treatment/State: G1 = G1 phase; M = metaphase; Oligo = HeLa oligonucleosomes; plunge = plunge frozen. Dose, in electrons / Å^2^. Pixel size is the unbinned pixel size as the specimen level. The K3 data were collected in super-resolution mode, with ½ the pixel size reported in the table. Pixel size therefore refers to the camera’s “bin ×1” pixel. Refined defocus (∆f) values are reported for defocus phase-contrast data. Nominal defoci are reported for Volta phase-contrast (VPP) data. ∆tilt = tilt increment. Fig = figures that show this dataset; those without a figure number were used for subtomogram averaging. Camera (Cam): FII = Falcon II, K3 = K3-GIF. thick = thickness, measured from the reconstructed cryotomogram. resid = alignment residual, in nanometers.

**Table S6. Subtomogram analysis of chromatin.**

|  | **Oligonucleosomes** | | **G1 phase lamella** | | | **Metaphase lamella** | | |
| --- | --- | --- | --- | --- | --- | --- | --- | --- |
|  | 0% DMSO | 9% DMSO | NCP Class 1 – 7 | | Di-NCP | NCP Class 1 – 8 | | Di-NCP |
| **Template matching** |  |  |  |  |  |  |  |  |
| Tomograms | 29 | 30 | 21 | | | 4 | | |
| Reference | Cylinder | Cylinder | Cylinder | | | Cylinder | | |
| Mask | Cylinder | Cylinder | Cylinder | | | Cylinder | | |
| **2-D classification** |  |  |  |  |  |  |  |  |
| Subtomograms | 1,136,628 | 1,280,772 |  | - |  |  | - |  |
| Classes (total) | 50 | 50 |  | - |  |  | - |  |
| Classes (kept) | 34 | 27 |  | - |  |  | - |  |
| T parameter | 2 | 2 |  | - |  |  | - |  |
| Mask (Å) | 110 | 110 |  | - |  |  | - |  |
| E-step (Å) | 25 | 25 |  | - |  |  | - |  |
| **3-D classification** |  |  |  |  |  |  |  |  |
| Subtomograms | 207,884 | 175,984 | 165,964 | | |  | 145,804 |  |
| Reference | Cylinder | Cylinder | Di-NCP STA (G1) | | | Di-NCP STA (Metaphase) | | |
| Mask, sphere (Å) | 110 | 110 | 240 | | | 240 | | |
| E-step (Å) | 20 | 20 | 20 | | | 20 | | |
| T parameter | 4 | 4 | 4 | | | 4 | | |
| Classes (total) | 30 | 30 | 50 | | | 50 | | |
| Classes (kept) | 2 | 3 | 3 | | 1 | 4 | | 1 |
| Symmetry imposed | C1 | C1 | C1 | | | C1 | | |
| **Refinement** |  |  |  |  |  |  |  |  |
| Subtomograms | 8,884 | 9,074 | 19,489 | | 4,199 | 17,224 | | 4,668 |
| Reference | Cylinder | Cylinder | NCP Class 1 – 7 STA  (G1) | | Di-NCP STA (G1) | NCP Class 1 – 8 STA (Metaphase) | | Di-NCP STA (Metaphase) |
| Mask, sphere (Å) | 110 | 110 | 240 | | 240 | 240 | | 240 |
| Symmetry imposed | C1 | C1 | C1 | | C1 | C1 | | C1 |
| Resolution (Å) |  |  |  |  |  |  |  |  |
| FSC = 0.5 | 19.0 | 19.5 | 30.6 – 33.7 | | 26.8 | 32.5 – 34.3 | | 26.3 |
| FSC = 0.143 | 16.7 | 18.1 | 24.3 – 27.4 | | 21.1 | 24.9 – 26.2 | | 21.3 |
| EMDB entry | EMD-37004 | EMD-37005 | EMD-36993 | | EMD-36992 | EMD-36999 | | EMD-36998 |

NCP = mononucleosomes, Di-NCP = stacked dinucleosomes, STA = subtomogram average. The values here reflect the final major round of 3-D classification (Figures S14D and S30D) and refinement (Figures S15 & S31, A & B).

**Table S7. Subtomogram analysis of chromatin – nucleosomes with gyre-proximal densities**

|  | **G1 phase lamella** | | | | | | |  | **Metaphase lamella** | | | | | |
| --- | --- | --- | --- | --- | --- | --- | --- | --- | --- | --- | --- | --- | --- | --- |
|  | Prox. 1 | Prox. 2 | | Prox. 3 | Prox. 4 | | Prox. 5 |  | Prox. 1 | Prox. 2 | | Prox. 3 | | Prox. 4 |
| **Template matching** |  | |  | | |  | |  |  | |  | |  | |
| Tomograms | 21 | | | | | | |  | 4 | | | | | |
| Reference | Cylinder | | | | | | |  | Cylinder | | | | | |
| Mask | Cylinder | | | | | | |  | Cylinder | | | | | |
| **2-D classification** |  | |  | | |  | |  |  | | | | | |
| Subtomograms |  | | - | | |  | |  |  | | - | |  | |
| Classes (total) |  | | - | | |  | |  |  | | - | |  | |
| Classes (kept) |  | | - | | |  | |  |  | | - | |  | |
| T parameter |  | | - | | |  | |  |  | | - | |  | |
| Mask (Å) |  | | - | | |  | |  |  | | - | |  | |
| E-step (Å) |  | | - | | |  | |  |  | | - | |  | |
| **3-D classification** |  | |  | | |  | |  |  | | | | | |
| Subtomograms | 165,964 | | | | | | |  |  | | 145,804 | |  | |
| Reference | Di-NCP STA (G1) | | | | | | |  | Di-NCP STA (Metaphase) | | | | | |
| Mask, sphere (Å) | 240 | | | | | | |  | 240 | | | | | |
| E-step (Å) | 20 | | | | | | |  | 20 | | | | | |
| T parameter | 4 | | | | | | |  | 4 | | | | | |
| Classes (total) | 50 | | | | | | |  | 50 | | | | | |
| Classes (kept) | 1 | | | | | | |  | 1 | | | | | |
| Symmetry imposed | C1 | | | | | | |  | C1 | | | | | |
| **Refinement** |  | |  | | |  | |  |  | | | | | |
| Subtomograms | 2,016 | 1,309 | | 1,947 | 2,126 | | 2,038 |  | 1,361 | 1,837 | | 1,725 | | 1,397 |
| Reference | Prox. 1 STA (G1) | Prox. 2 STA (G1) | | Prox. 3 STA (G1) | Prox. 4 STA (G1) | | Prox. 5 STA (G1) |  | Prox. 1 STA (Metaphase) | Prox. 2 STA (Metaphase) | | Prox. 3 STA (Metaphase) | | Prox. 4 STA (Metaphase) |
| Mask, sphere (Å) | 240 | 240 | | 240 | 240 | | 240 |  | 240 | 240 | | 240 | | 240 |
| Symmetry imposed | C1 | C1 | | C1 | C1 | | C1 |  | C1 | C1 | | C1 | | C1 |
| Resolution (Å) |  | |  | | |  | |  |  | |  | |  | |
| FSC = 0.5 | 30.6 | 33.4 | | 31.3 | 33.0 | | 33.7 |  | 32.5 | 32.6 | | 33.4 | | 34.3 |
| FSC = 0.143 | 24.3 | 25.5 | | 25.3 | 27.4 | | 26.4 |  | 25.6 | 24.9 | | 25.9 | | 26.2 |
| EMDB entry | EMD-36994 | | | | | | |  | EMD-37000 | | | | | |

Prox. = Nucleosomes with gyre-proximal density, STA = subtomogram average. The values here reflect the final major round of 3-D classification (Figures S14D and S30D) and refinement (Figures S15C and S31C).

**Table S8. Subtomogram analysis of megacomplexes.**

|  | **G1 phase lamella** | | | **Metaphase lamella** |
| --- | --- | --- | --- | --- |
|  | Ribosome (Cytoplasm only) | Ribosome  (Full volume) | Preribosome  (Full volume) | Ribosome |
| **Template matching** |  |  |  |  |
| Tomograms | 11 | 21 | | 4 |
| Reference | Sphere | Sphere | | Sphere |
| Mask | Sphere | Sphere | | Sphere |
| **2-D classification** |  |  |  |  |
| Subtomograms | - | 192,309 | | - |
| Classes (total) | - | 100 | | - |
| Classes (kept) | - | 15 | | - |
| T parameter | - | 2 | | - |
| Mask (Å) | - | 300 | | - |
| E-step (Å) | - | 20 | | - |
| **3-D classification** |  |  |  |  |
| Subtomograms | 11,972 | 25,886 | | 48,849 |
| Reference | Sphere | Ribosome STA | | Ribosome STA |
| Mask, sphere (Å) | 300 | 300 | | 300 |
| E-step (Å) | 50 | 50 | | 50 |
| T parameter | 4 | 4 | | 4 |
| Classes (total) | 30 | 30 | | 50 |
| Classes (kept) | 1 | 1 | 1 | 1 |
| Symmetry imposed | C1 | C1 | | C1 |
| **Refinement** |  |  |  |  |
| Subtomograms | 684 | 702 | 635 | 332 |
| Reference | Ribosome STA | Ribosome STA | Preribosome STA | Ribosome STA |
| Mask, sphere (Å) | 300 | 300 | 300 | 300 |
| Symmetry imposed | C1 | C1 | C1 | C1 |
| Resolution (Å) |  |  |  |  |
| FSC = 0.5 | 31.7 | 33.4 | 33.3 | 36.8 |
| FSC = 0.143 | 24.7 | 24.7 | 25.4 | 26.9 |
| EMDB entry | EMD-37002 | | EMD-37001 | EMD-37003 |

STA = subtomogram average.

**Table S9. Subtomogram analysis of G1 chromatin with 9% glycerol or without cryoprotectant.**

|  | **G1 with 9% glycerol cryoprotectant** | |  | **G1 without cryoprotectant** |
| --- | --- | --- | --- | --- |
|  | NCP Class 1 – 6 | Prox. |  | NCP Class 1 – 9 |
| **Template matching** |  |  |  |  |
| Tomograms | 5 | |  | 9 |
| Reference | Cylinder | |  | Cylinder |
| Mask | Cylinder | |  | Cylinder |
| **2-D classification** |  | |  |  |
| Subtomograms | - | |  | - |
| Classes (total) | - | |  | - |
| Classes (kept) | - | |  | - |
| T parameter | - | |  | - |
| Mask (Å) | - | |  | - |
| E-step (Å) | - | |  | - |
| **3-D classification** |  |  |  |  |
| Subtomograms | 187,245 | |  | 175,229 |
| Reference | NCP STA (G1) | |  | NCP STA (Metaphase) |
| Mask, sphere (Å) | 240 | |  | 240 |
| E-step (Å) | 20 | |  | 20 |
| T parameter | 4 | |  | 4 |
| Classes (total) | 50 | |  | 50 |
| Classes (kept) | 3 | 1 |  | 4 |
| Symmetry imposed | C1 | |  | C1 |
| **Refinement** |  |  |  |  |
| Subtomograms | 12,607 | 1,670 |  | 17,312 |
| Reference | NCP Class 1 – 6 STA | Prox. STA |  | NCP Class 1 – 9 STA |
| Mask, sphere (Å) | 240 | 240 |  | 240 |
| Symmetry imposed | C1 | C1 |  | C1 |
| Resolution (Å) |  |  |  |  |
| FSC = 0.5 | 28.8 – 34.0 | 31.4 |  | 29.4 – 55.7 |
| FSC = 0.143 | 24.3 – 28.6 | 26.0 |  | 25.9 – 31.3 |
| EMDB entry | EMD-xxxxx | EMD-xxxxx |  | EMD-xxxxx |

NCP = mononucleosomes, Prox. = Nucleosomes with gyre-proximal density. The values here reflect the final major round of 3-D classification and refinement shown in Figures S18 & S19 (9% glycerol) and Figures S21 & S22 (no cryoprotectant).
